# Structure of a Tc holotoxin pore provides insights into the translocation mechanism

**DOI:** 10.1101/590430

**Authors:** Daniel Roderer, Oliver Hofnagel, Roland Benz, Stefan Raunser

**Affiliations:** Department of Structural Biochemistry, Max Planck Institute of Molecular Physiology, Otto-Hahn-Str. 11, 44227 Dortmund, Germany; Department of Life Sciences and Chemistry, Jacobs University Bremen, Campusring 1, 28759 Bremen, Germany

## Abstract

Tc toxins are modular toxin systems that are composed of a pentameric membrane translocator (TcA) and a cocoon (TcB and TcC) encapsulating the toxic enzyme. Binding of Tcs to target cells and a pH shift trigger the conformational transition from the soluble prepore state to the membrane-embedded pore. Subsequently, the toxic enzyme is translocated and released into the cytoplasm. Here, we show in atomic detail an assembled Tc toxin complex from *P. luminescens* in the membrane. We find that the five TcA protomers conformationally adapt to fit around the cocoon during prepore-to-pore transition. The architecture of the Tc toxin complex also allows TcB-TcC to bind to an already membrane-embedded TcA pore to form a holotoxin. Mammalian lipids with zwitterionic head groups are preferred over other lipids for Tc toxin integration. The translocated toxic enzyme, which can be partially visualized, transiently interacts with alternating negative charges and hydrophobic stretches.

## Introduction

*Photorhabdus* toxin complexes (Tc toxins) are large, heterotrimeric protein complexes that are produced by many insect and human pathogenic bacteria (Waterfield et al., 2001) (ffrench-Constant and Waterfield, 2005). Tcs were originally discovered in *Photorhabdus luminescens* (Bowen et al., 1998). Later, gene loci of Tc toxins were also found in *Xenorhabdus nematophila* (ffrench-Constant and Bowen, 2000; Sergeant et al., 2003), and human pathogens, such as *Photorhabdus asymbiotica* (Gerrard et al., 2004) and different species of *Yersinia* (Tennant et al., 2005; Waterfield et al., 2007). While the entomopathogenic toxins are potential biopesticides and therefore the focus of crop protection research, understanding the mechanism of action of Tc toxins of human pathogens is medically relevant.

A Tc toxin is typically made up of three components, TcA, TcB and TcC. *P. luminescens* have different TcA, TcB and TcC components (Duchaud et al., 2003; ffrench-Constant et al., 2000), enabling them to form various holotoxins that result in the translocation of different toxic payloads (Lang et al., 2010). The modularity of Tc toxins in insect pathogens may provide an alternative to Bt toxins for green biotechnology, overcoming the problem of rising resistance in insects (Tabashnik et al., 2013).

TcA is a ∼ 1.4 MDa pentameric protein that acts as membrane permeation and protein translocation device. It is 18 nm wide and 24 nm long and is shaped like a bell. The α-helical translocation channel at the center of TcA is hydrophobic at its tip and shielded by a large shell (Gatsogiannis et al., 2013). Receptor-binding domains at the periphery of the shell likely mediate the initial receptor-toxin interaction on target cells (Meusch et al., 2014). A shift to higher or lower pH opens an electrostatic lock at the bottom of the shell, triggering its structural rearrangement and the release of the shielded translocation channel. Membrane permeation of the channel is driven by a proline-rich linker that connects it with the shell. The linker condenses from a stretched and unfolded conformation in the prepore to a partially helically folded, compacted conformation in the pore, resulting in a shifting-out movement of the channel from the shell (Gatsogiannis et al., 2016). Once inside the membrane, the channel opens and releases the toxic enzyme into the cytosol (Gatsogiannis et al., 2016).

TcB and TcC form a ∼ 250 kDa cocoon which is mainly composed of RHS (rearrangement hotspot) repeats. The C-terminal end of the cocoon protrudes inwards and forms an aspartyl autoprotease domain that cleaves off the C-terminal hypervariable region (HVR) of TcC (Meusch et al., 2014). This generates a toxic enzyme of ∼ 30 kDa that resides inside the cocoon. In contrast to the highly conserved N-terminal region of TcC, the sequence of the C-terminus of TcC is not conserved and therefore called HVR (Lang et al., 2010). The various HVRs are toxic enzymes that have either different functions or targets. For example, while the toxic enzymes of TccC3 and TccC5 from *P. luminescens* act both as ADP-ribosyltransferases, they modify different proteins; TccC3 ADP-ribosylates actin and TccC5 ADP-ribosylates Rho GTPases (Lang et al., 2010). The hydrophobic inside of the cocoon creates an environment unfavorable for protein folding, most likely resulting in an unfolded conformation of the toxic enzyme (Busby et al., 2013; Meusch et al., 2014).

The TcB-TcC cocoon binds through a distorted β-propeller that closes it at the bottom to the funnel-shaped end of TcA, which results in the ABC holotoxin complex (Meusch et al., 2014). During this process, parts of the β-propeller completely unfold and refold into an alternative conformation, resulting in the opening of the cocoon and release of the toxic enzyme into the channel with the C-terminus first (Gatsogiannis et al., 2018).

Several structures of TcA in its prepore and pore state as well as structures of the ABC holotoxin complex enabled us to understand the major steps in the mechanism of Tc toxin action (Gatsogiannis et al., 2016; 2018; Meusch et al., 2014). However, a structure of the ABC complex in its pore state is missing, hampering our understanding of how the complete complex enters the membrane and translocates its toxic payload.

Here, we present a near-atomic cryo-EM structure of an ABC holotoxin from *P. luminescens* embedded in lipid nanodiscs, which reveals that the shell domain of TcA forms a basin, in which the TcB-TcC complex resides. The cocoon interacts directly with the α-helical shell domains of two out of five TcA subunits, which have to adapt their conformation slightly to fit around the TcB-TcC cocoon. In general, all upper TcA domains are better resolved than in the TcA complex alone, indicating that they are stabilized by the cocoon (Gatsogiannis et al., 2016). We show that the assembled ABC complex can directly enter the membrane. Lipids with zwitterionic head groups and mammalian lipids are preferred over lipids with negatively charged head groups and bacterial lipids. Interestingly, the stable TcA-TcB interface is not affected by the prepore-to-pore transition of TcA, and TcB-TcC can also bind to the already membrane-embedded TcA pore to form a holotoxin. Furthermore, the structure reveals that the translocated toxic enzyme transiently interacts with alternating hydrophobic and negatively charged regions in the translocation channel.

## Results and Discussion

### Structure of the ABC holotoxin complex in nanodiscs

In order to obtain an ABC complex where the toxic enzyme is still residing inside the translocation channel, we assembled the ABC holotoxin from the individually purified toxin components with proteolytically inactive TcC (TcdA1, TcdB2-TccC3(D651A) *from P. luminescens*, referred to collectively as ABC(D651A)) (Gatsogiannis et al., 2018; Meusch et al., 2014), which is, in contrast to the wild-type ABC holotoxin (ABC(WT)), non-toxic to HEK 293T cells (Supplementary Figure 1a). We induced pore formation and simultaneous reconstitution in lipid nanodiscs by dialyzing a mixture of ABC(D651A) and nanodiscs against a buffer at pH 11 (Supplementary Figure 1b-c). We then solved the structure of the complex by cryo-EM and single particle analysis using SPHIRE (Moriya et al., 2017) at an average resolution of 3.4 Å (Figure 1a,b, Supplementary Figure 2, Supplementary Table 1, Supplementary Movie 1). To improve the local resolution of the TcB-TcC cocoon, we shifted the center towards these components during image processing (Methods, Supplementary Figure 3).

**Figure 1:**
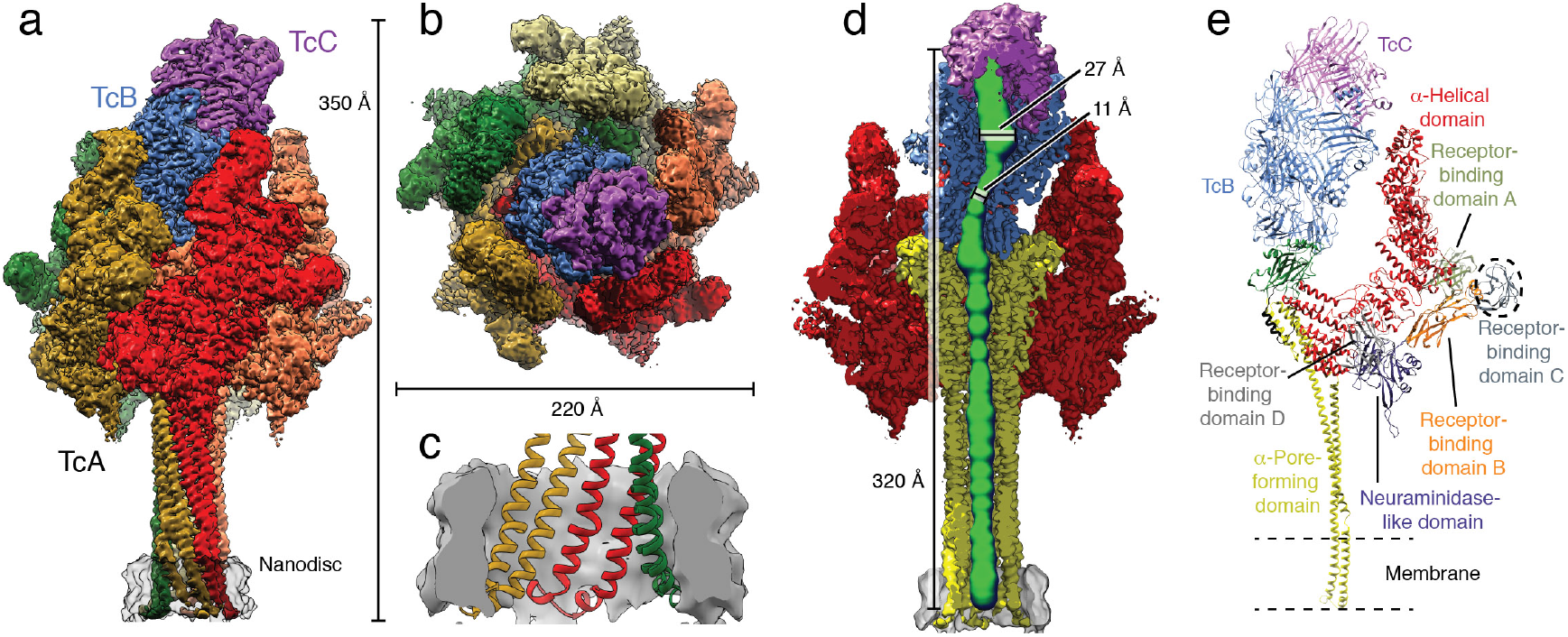
Structure of the Tc holotoxin in a lipid nanodisc. **a,b**: Cryo-EM map of ABC(D651A) in side view (**a**) and top view (**b**). The individual protomers of TcA (A-E), TcB and TcC are shown in different colors. The lipid nanodisc is shown in transparent gray. **c**: Longitudinal section through the nanodisc region. The density of the nanodisc is shown in gray, and the molecular model of the TcA transmembrane domains is colored by protomer. **d**: Longitudinal section through the cryo-EM structure displaying the central channel. The densities corresponding to the TcA outer shell, the TcA channel, TcB, TcC and the nanodisc are shown in red, yellow, blue, violet and gray, respectively. The lumen of the translocation channel was determined in ChExVis (Masood et al., 2015) and is shown in green. Diameters inside the cocoon and at the constriction site are indicated. **e**: Molecular models of one TcA protomer, TcB and TcC in the context of the holotoxin. TcA is colored by domains. Receptor-binding domain C (dashed circle) is not resolved in the cryo-EM structure. TcB is colored blue, and TcC is colored violet.

The high quality of the map allowed us to build a model of 91% of the pore state of ABC(D651A). The final model contains residues 89-2516 of TcA, residues 1 – 1471 of TcB and residues 1 – 683 of TcC with some loops missing. The only parts of the complex that could not be resolved are the receptor-binding domain (RBD) C of TcA (residues 1382-1491) and the ADP-ribosyltransferase of TcC (residues 684-960), indicating that these regions are highly flexible (Figure 1e).

The structure of TcA in the 1.7 MDa ABC pore complex resembles very much the structure of the isolated TcA subunit embedded in lipid nanodiscs (pdb ID 5LKI) (Gatsogiannis et al., 2016) (Supplementary Figure 4a,b). The translocation channel is open and extends all the way from the TcB-TcC cocoon to the transmembrane region in the nanodisc (Figure 1c,d).

The interface between the β-propeller of TcB and the TcB-binding domain of TcA is practically identical to that observed in the ABC prepore state (pdb ID 6H6F) (Supplementary Figure 4c,d) (Gatsogiannis et al., 2018). This and the rigidity of the rest of the cocoon result in the same relative position of the TcB-TcC cocoon to TcA, which sits at an angle of 32° on top of TcA (Supplementary Figure 4c). As a consequence, one of the five α-helical domains of the shell that form a tight basin around TcB-TcC is shifted outwards, which results in a break of the C5 symmetry of TcA at this position (Figure 2a,d, Supplementary Movie 2). The shift of TcA-A relative to the other TcA subunits is most pronounced at the uppermost part of the α-helical shell (residues 645-797) (Figure 2b-d). The other domains of TcA-A, such as the RBDs, the neuraminidase-like domain, and the α-helical pore-forming domain remain unchanged (Figure 2c), indicating that the conformational changes at the tip of the shell are not transmitted to the rest of the protein.

**Figure 2:**
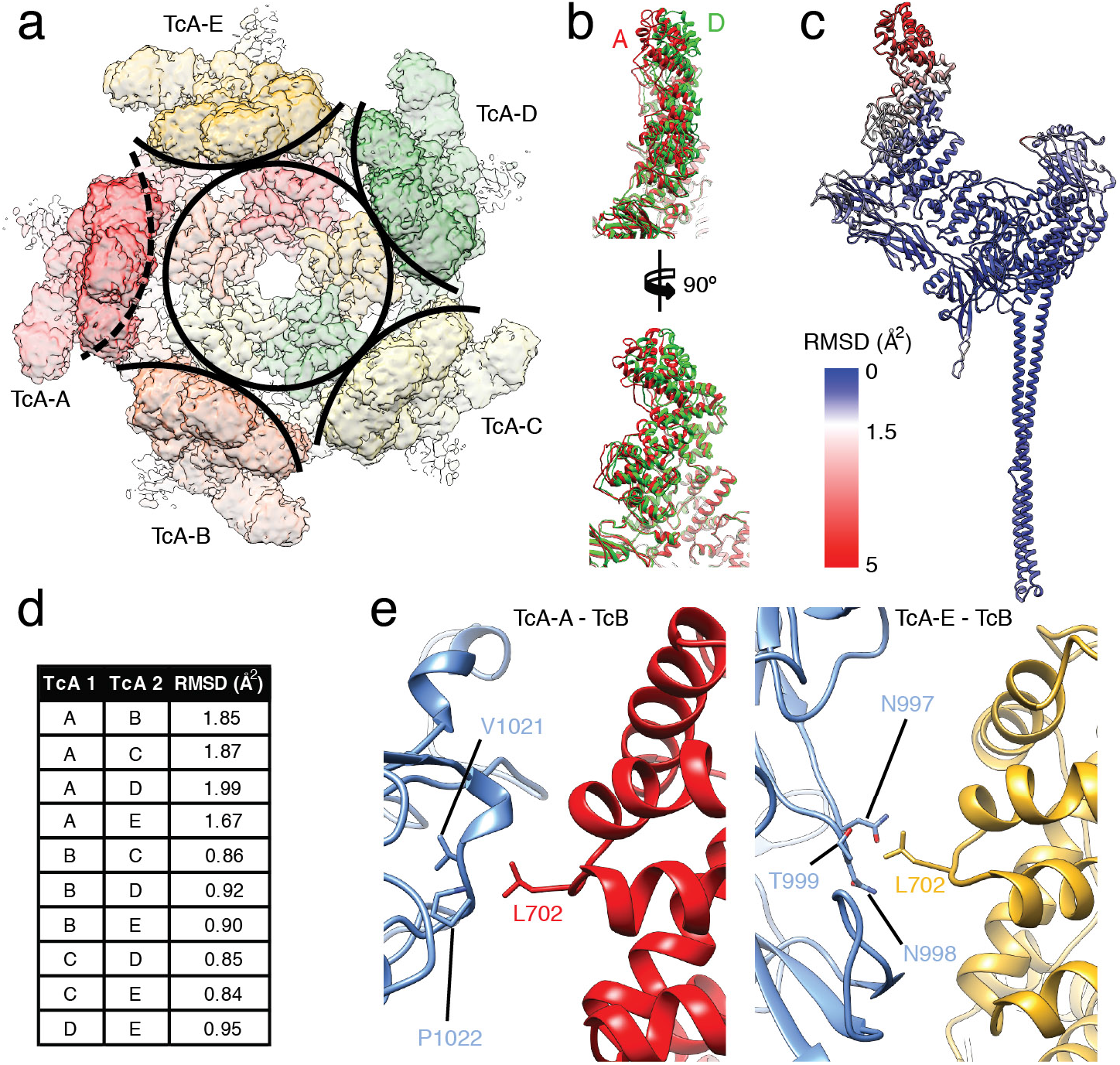
Interaction of the TcB-TcC cocoon with the TcA shell. **a**: Top view of the density map corresponding to TcA in the context of ABC(D651A). A circle around the center of the TcA channel and arcs around the outer shell of individual TcA protomers indicate the disruption of C5 symmetry by TcA-A (dashed arc). **b**: Overlay of the upper parts (residues 290 – 980) of TcA protomers A (red) and D (green). Front view (top) and side view (bottom) show a gradual increase of displacement with further distance to the center of TcA. **c**: RMSD of C*_α_* atoms of the individual TcA protomers, plotted on TcA-A. RMSD values (in Å^2^) increase from blue to red. **d**: Backbone RMSDs between the individual TcA chains, calculated in Chimera. **e**: Interaction of the cocoon (TcB, blue) with two out of five TcA protomers. Left panel: Interaction of L702 of TcA-A (red) with V1021 and P1022 of TcB. Right panel: Interaction of L702 of TcA-E (ochre) with N997, N998 and T999 of TcB.

The TcB-TcC cocoon directly interacts with the upper α-helical shell domain of two TcA chains. L702 of the displaced TcA protomer (TcA-A) and the same residue of the adjacent TcA-E interact with two hydrophobic patches at the surface of the cocoon (Figure 2e). Although there are no close contacts between the cocoon and the other three TcA protomers, the entire upper α-helical shell of the pentamer is stabilized in the ABC pore, likely due to a decreased level of freedom of these domains in the holotoxin. Thus, in contrast to the structure of the TcA pore in the absence of the cocoon (Gatsogiannis et al., 2016), the complete upper part of the outer shell is resolved (Supplementary Figure 2e, Supplementary Figure 4a,b).

Unexpectedly, the density corresponding to the ADP-ribosyltransferase does not reach the lower end of the translocation channel or is translocated across the membrane (Figure 3b, Supplementary Movie 3). Instead, it resides in the cocoon and only in the very upper part of the TcA translocation channel (Figure 3b-e), comparable to the ADP-ribosyltransferase in the prepore form of the cleavage-deficient ABC(D651A) (Gatsogiannis et al., 2018). This suggests that either the covalently linked toxic enzyme is locked in the cocoon and does not have the necessary flexibility to be further translocated, or a missing driving force such as a pH gradient could be responsible for the halted translocation.

**Figure 3:**
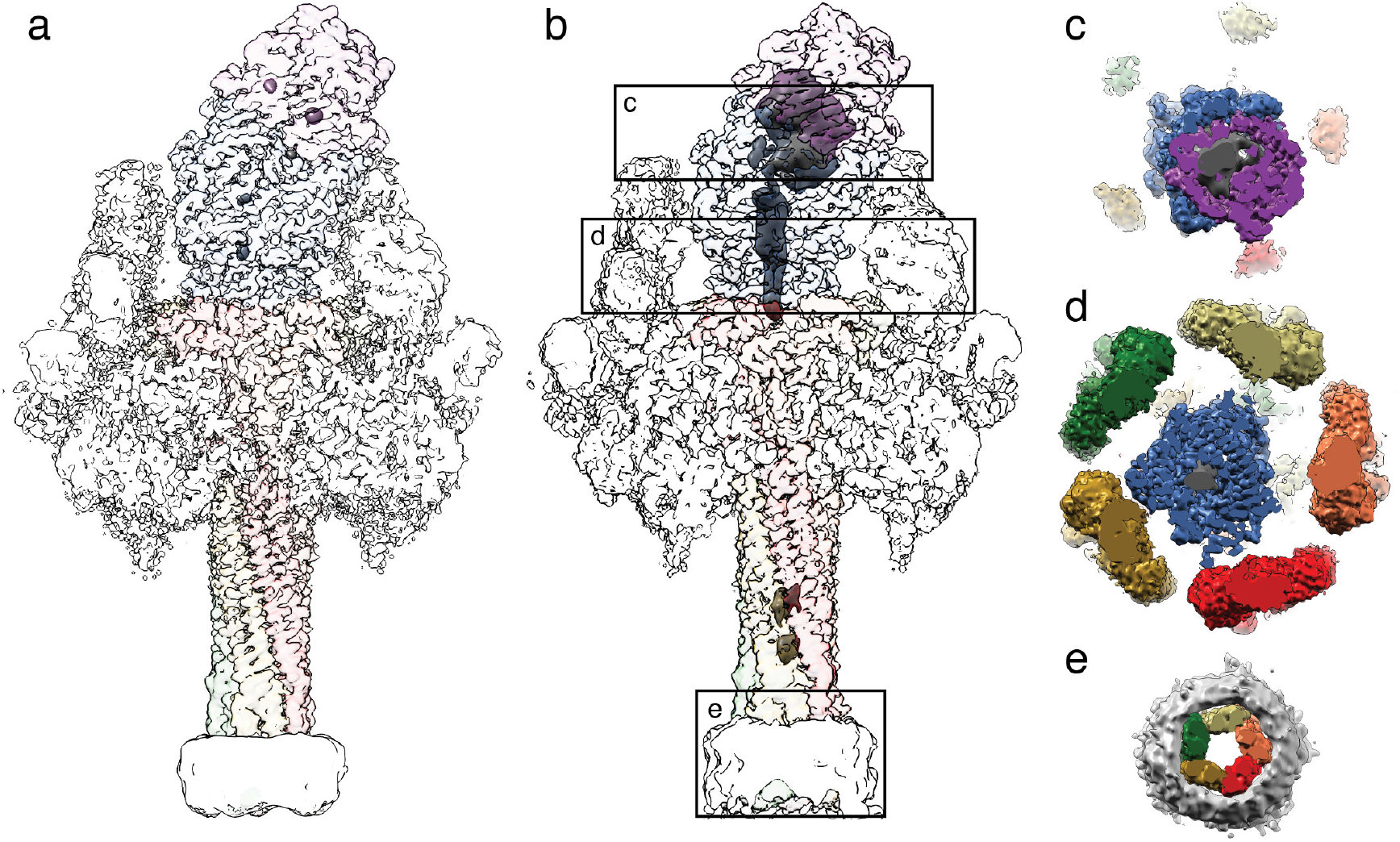
ADP-ribosyltransferase in the ABC(D651A) pore. **a,b**: Reconstruction of ABC(WT) (**a**) and ABC(D651A) (**b**) with transparent surface. Density, in which no atomic model was fitted, and the nanodisc was filtered to 10 Å and is shown at a lower threshold. **c,d,e**: cross-sections through ABC(D651A) at the cocoon, the TcB translocation channel and the transmembrane region, as indicated in panel b. Continuous density of the ADP-ribosyltransferase is apparent in the cocoon (**c**) and in the narrow passage of TcB below the constriction site (**d**), but not in the transmembrane region embedded in the nanodisc (**e**).

To find out whether the latter is true, we solved the structure of the ABC(WT) embedded in lipid nanodiscs at an average resolution of 3.4 Å (Supplementary Figure 5). If the translocation of the toxic enzyme is indeed dependent on an additional driving force, the protein should as well reside inside the cocoon and the TcA channel, comparable to the ABC(D651A). However, while the rest of the cryo-EM map did not differ from that of ABC(D651A) in its pore state, we did not find any density corresponding to the ADP-ribosyltransferase (Figure 3a, Supplementary Movie 3). This demonstrates that the ADP-ribosyltransferase is translocated and released from the holotoxin. Since we reconstituted the protein *in vitro* in the absence of other proteins, we conclude that no additional factors like molecular chaperones or driving forces, such as a pH gradient are necessary to translocate the ADP-ribosyltransferase. In addition, translocation of the ADP-ribosyltransferase requires the protein to be cleaved and to be flexible enough to move inside the TcB-TcC cocoon. Otherwise its translocation can only be initiated but not continued.

### ADP-ribosyltransferase inside the cocoon

The entire cocoon is resolved better than 4 Å in the ABC(D651A) pore complex, including the aspartyl protease cleavage site of TcC. As in the prepore complex, the density of the ADP-ribosyltransferase inside the cocoon and in the very upper part of the TcA translocation channel appears only at lower thresholds and no secondary structure elements are recognizable (Figure 3b). We estimated the 3D variance of the reconstruction in SPHIRE (Moriya et al., 2017). Expectedly, the highest variance is found inside the cocoon and also at the upper domains of TcA, indicating a high flexibility and disorder in these protein regions (Supplementary Figure 6a-c). 3D sorting focused on the inside of TcB-TcC did also not improve the quality of the density.

Nevertheless, due to the overall improved local resolution in the cocoon after shifting the center of reconstruction (Supplementary Figure 3e), we could identify the positions of the first five residues of TcC after the aspartyl protease cleavage site (M679 – A683). The residues form a loop that keeps the cleavage site between the catalytic aspartates (Figure 4a, Supplementary Figure 6d,e, Supplementary Movie 4). G677 and P680 are the only conserved residues around the autoproteolytic cleavage site (Figure 4a, b). To identify whether these residues are part of the recognition sequence defining the specificity of the protease, we mutated both residues to alanines. Surprisingly, P680A does not affect autoproteolysis and the mutation of G677 to alanine results in only ∼ 40% of the protein being uncleaved (Figure 4c). Thus, although these residues are conserved in RHS repeat proteins that function as toxins (not in teneurins (Jackson et al., 2018; Li et al., 2018)), neither of them is absolutely necessary for the activity of the aspartyl protease, indicating that there is no specific recognition sequence required. Similarly, aspartyl proteases of the digestive system, such as pepsin, that cleave in a rather unspecific manner adjacent to large hydrophobic and aromatic residues, act independently of the residues two positions upstream and downstream of the cleavage site (Athaudaa and Takahashia, 2002).

**Figure 4:**
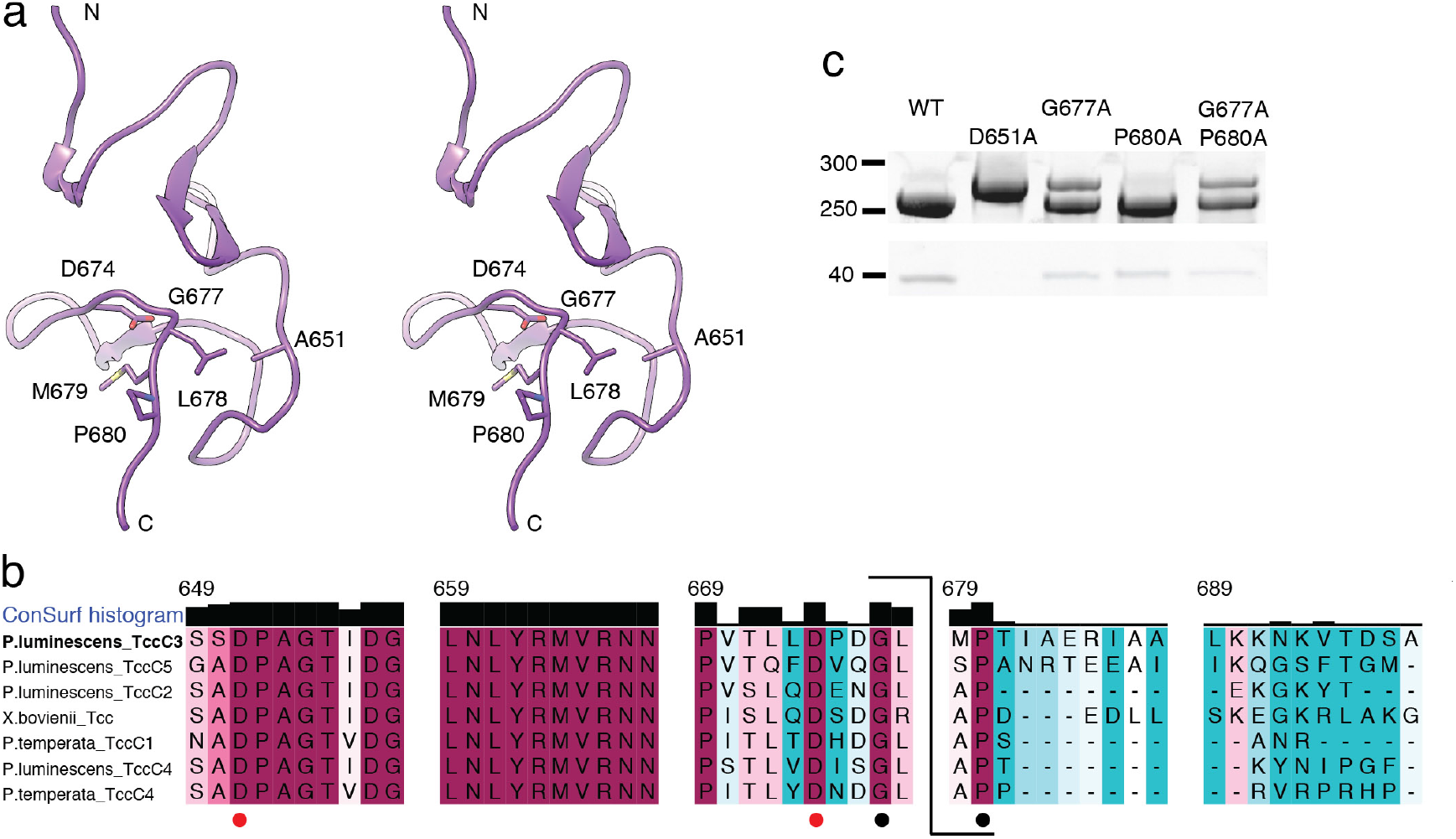
Structure and conservation of the aspartyl autoprotease domain of TcC. **a**: Stereo view of the aspartyl protease domain (residues 618 – 678) with the N-terminus of the ADP-ribosyltransferase in the TcB-TcC cocoon. The toxin is cleaved between L678 and M679. The residues forming the aspartyl protease cleavage site (D674 and A651, which is D651 in WT) and the conserved G677 and P680 are indicated. Representations that include the cryo-EM map are shown in Supplementary Figure 6d,e. **b**: Sequence alignment of the C-terminal part of the aspartyl autoprotease site and the N-terminal region of the toxic enzyme, colored from minimum (cyan) to maximum (magenta) conservation. The cleavage site is indicated (bracket). D651 and D674 of the aspartyl protease domain are marked with red dots, and the conserved G677 and P680 flanking the cleavage site are marked with black dots. **c**: Cleavage of the toxin by different mutations in the TcC cleavage site, analyzed by SDS-PAGE. TcC variant D651A does not cleave the toxin, and variant G677A shows impaired cleavage, as well as the double mutant G677A-P680A. Mutation P680A does not impair toxin cleavage significantly.

Since we could not determine the structure of the ADP-ribosyltransferase inside the TcB-TcC cocoon, we performed cross-linking mass spectrometry (XL-MS) using the 12 Å long cross-linker disuccinimidyl dibutyric urea (DSBU) to obtain information on the orientation of the protein inside the TcB-TcC(WT) cocoon (Supplementary Table 2). We selectively screened for crosslinks between the TcB-TcC cocoon and the ADP-ribosyltransferase and identified several of them at different positions of the molecules (Figure 5a). For example, K107 of the ADP-ribosyltransferase was crosslinked to S139, Y919, Y921 and K2104 of the cocoon (Figure 5a,b). The same crosslinks were found for TcB-TcC(D651A) with non-cleaved ADP-ribosyltransferase (Supplementary Figure 6f,g). These three positions are distributed over the entire inner surface of the TcB-TcC cocoon. In TcB-TcC(D651A) we found another prominent set of crosslinks between three lysines in the central part of the ADP-ribosyltransferase (K175, K178 and K185) and the residues S139, Y921, K1521 and K1761 of the cocoon. Again they are distributed over the entire lumen of the cocoon, indicating that the toxic enzyme does not have a fixed position inside the cocoon (Supplementary Figure 6f,h).

**Figure 5:**
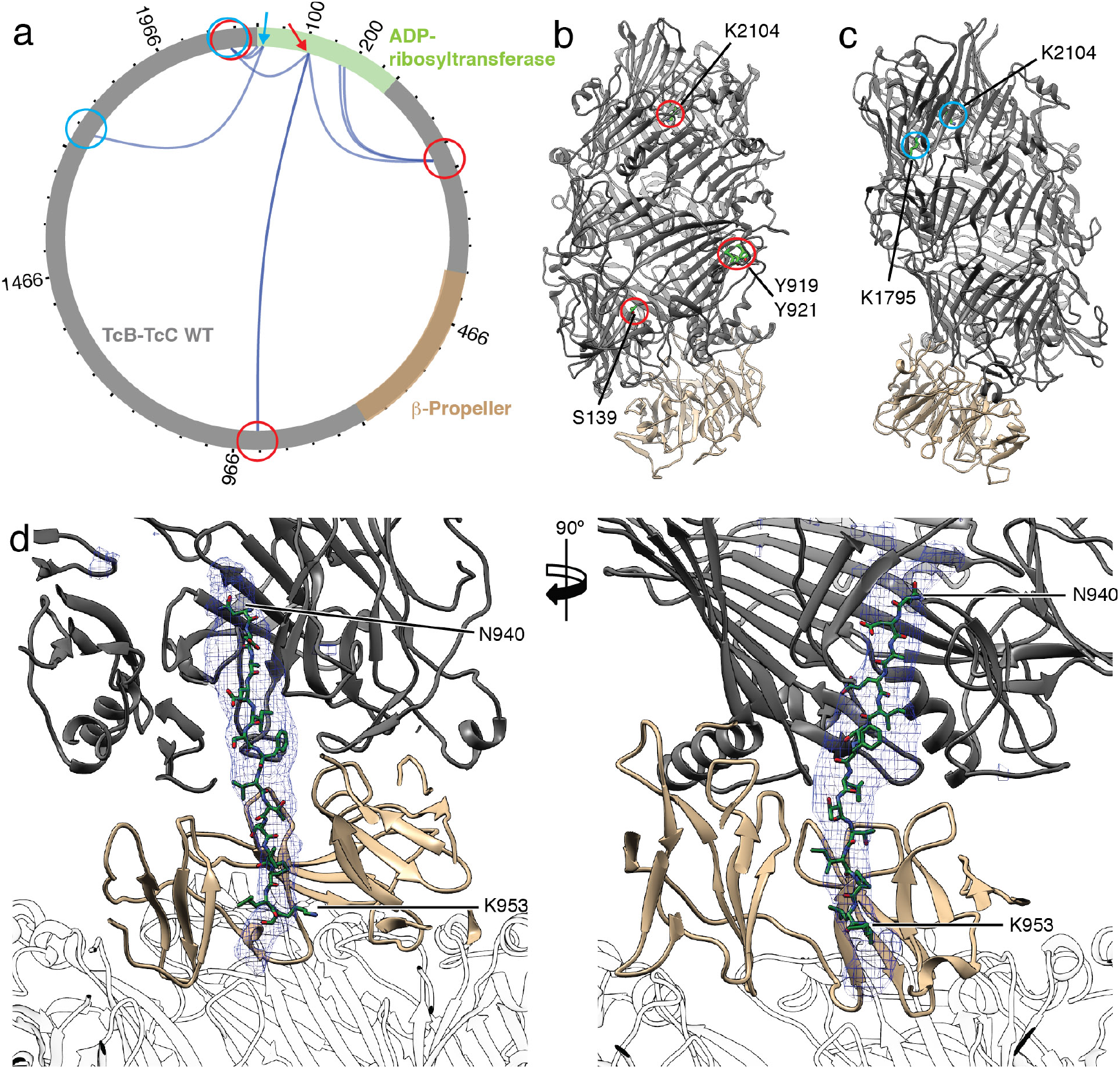
XL-MS of the toxic enzyme inside the TcB-TcC WT cocoon and atomic model of the ADP-ribosyltransferase in the narrow passage of TcB. **a**: Visualization of crosslinks (dark blue curves) between the TcB-TcC cocoon (gray, β-propeller in sand) and the toxic enzyme (green). Two individual positions (K107 and K12) and their crosslinks to the TcB-TcC cocoon are indicated with red and blue arrows and circles, respectively. All crosslinks with a score of 50 and higher are shown. The plot was created using xVis (Grimm et al., 2015). A complete list of all crosslinks is shown in Supplementary Table 2. **b,c**: The TcB-TcC cocoon (pdb ID 4O9X) with all amino acids that undergo crosslinks with K107 indicated and highlighted with red circles (**b**) and crosslinks with K12 indicated and highlighted with blue circles (**c**). The cocoon is colored gray, the β-propeller (residues 376-691) is colored sand. Residues are numbered according to pbd ID 6H6E. d: Atomic model of the ADP-ribosyltransferase (N940 – K953 of TcC) in the narrow passage of TcB. The density map corresponding to the ADP-ribosyltransferase is shown as blue mesh. TcA is shown transparent.

However, since K12, which is positioned close to the N-terminus of the ADP-ribosyltransferase, crosslinks to two prominent lysines close to the aspartyl protease site of the cocoon (K1795 and K2104), the N-terminus of the ADP-ribosyltransferase in TcB-TcC(WT) seems not to change its position after cleavage (Figure 5a,c).

Taken together, the XL-MS of TcB-TcC(WT) and TcB-TcC(D651A) indicate that the N-terminus of the ADP-ribosyltransferase resides close to its cleavage site whereas the rest of the protein takes random orientations inside the cocoon suggesting that the enzyme is in an unfolded or semi-unfolded state.

### The C-terminus of the ADP-ribosyltransferase inside the translocation channel

Before entering the TcA translocation channel, the toxic enzyme has to pass a narrow passage at the lower part of the cocoon (Gatsogiannis et al., 2018) (Figure 1d, Supplementary Figure 8a-c). Although the density corresponding to the ADP-ribosyltransferase in this region was better resolved in the pore structure of ABC(D651A) than in our previous prepore structure (Gatsogiannis et al., 2018), it was still difficult to build an atomic model of the protein. Since it is obvious that the protein enters the translocation channel with the C-terminus first, we tested the optimal fit of three different C-terminal fragments of TcC (I932 – S945, N940 – K953, and L947 – R960) and found that the peptide N940-K953 fitted the best into the density (Figure 5d, Supplementary Figure 7a,b, Supplementary Movie 5). Although secondary structure predictions suggest the presence of a short β-strand at the center of the peptide (Supplementary Figure 7a), it fits only in the density in its stretched, unfolded conformation, indicating that the protein passes the constriction site in its unfolded state similar to translocated substrates in HSP104 (Gates et al., 2017) or AAA+ proteases (Puchades et al., 2017).

The structure of the C-terminus of the ADP-ribosyltransferase inside the narrow passage of TcB reveals that along its translocation pathway, the enzyme passes first through a conserved negatively charged constriction site formed by D34, N60, D73 and E102 (Gatsogiannis et al., 2018), and then through a conserved hydrophobic stretch (F741, I743, W771 and F778). Afterwards, it passes through a negatively charged ring and finally a hydrophobic ring inside the β–propeller (Figure 6a-c, Supplementary Figure 8d,g). Interestingly, the protein does not locate at the axis of the channel but adheres to the channel wall (Figure 6d,e). As expected, the interactions are weak but polar and hydrophobic residues of the ADP-ribosyltransferase are stabilized by interacting with hydrophilic and hydrophobic stretches of the channel, respectively (Figure 6c-e).

**Figure 6:**
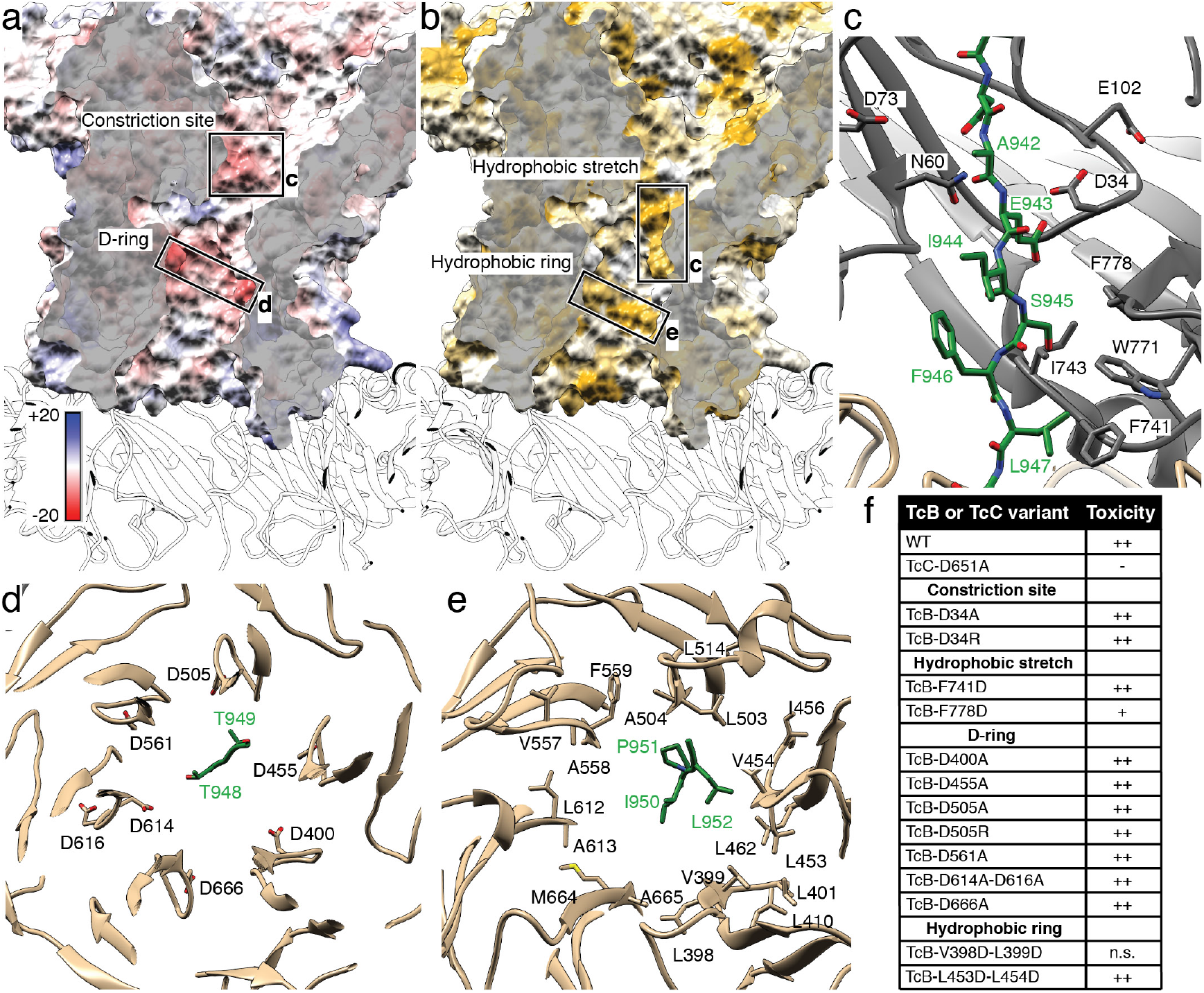
Alternating hydrophobic and negatively charged stretches in the narrow passage of TcB in ABC(D651A). **a**: Surface representation of the narrow passage of TcB, colored according to the Coulomb potential (kcal mol^−1^ e^−1^) at pH 7.0. The negatively charged constriction site at the channel entrance and a negatively charged ring (D-ring) in the center of the channel are highlighted in dashed boxes. Translocation direction is from top to bottom. **b**: The same view as in panel a, colored according to hydrophobicity as described in Hessa *et al*., 2005 (Hessa et al., 2005). Hydrophobic regions are colored ochre, non-hydrophobic regions are colored white. A hydrophobic stretch at the upper part of the channel and a hydrophobic ring close to the TcB-TcA interface are highlighted. **c**: Section of the narrow passage of TcB showing the hydrophobic stretch made up of L739, F741, I743, W771 and F778. D34, N60, D73 and E102 form the negatively charged constriction site at the entrance of the TcB channel. The translocating ADP-ribosyltransferase (A942 – L947 of TcC) is shown in green. **d**: The D-ring of TcB is formed by D400, D455, D505, D561, D614, D616 and D666. T948 and T949 of TcC interact with the D-ring. **e**: The hydrophobic ring is formed by L398, V399, L401, L410, L453, V454, I456, L462, L503, A504, L514, V557, A558, F559, L612, A613, M664 and A665. I950-L952 of TcC are at the height of the hydrophobic ring. **f**: Toxicity of TcB-TcC variants against HEK293-T cells in the context of the holotoxin. n.s.: non-soluble expression of the respective variant.

The negatively charged ring inside the β–propeller is formed by seven conserved aspartate residues (D400, D455, D505, D561, D614, D616 and D666) (Figure 6a,d, Supplementary Figure 8e,h). We therefore call it the D-ring. It interacts with the polar residues T948 and T949 of the ADP-ribosyltransferase (Figure 6d).

The hydrophobic ring, which forms the central section of the β–propeller is made up of many small and large hydrophobic residues facing the channel lumen (Figure 6b,e). They are either strictly conserved or replaced by nearly equally sized hydrophobic side chains (Supplementary Figure 8f,h). The ring interacts transiently with three hydrophobic residues of the ADP-ribosyltransferase (I950, P951, L952).

To demonstrate the role of these striking features in TcB-TcC, we mutated several key residues to change their property from hydrophobic to hydrophilic or negatively charged to either neutral or positively charged and probed the toxicity of the variants on HEK 293T cells (Supplementary Figure 9). Surprisingly, the insertion of charged residues into the hydrophobic stretch has no or only minor impairing effects (Figure 6f, Supplementary Figure 9). Similarly, mutations of L453 and L454 to aspartate in the hydrophobic ring have no effect on toxicity and mutations of L398 and L399 to alanine cause insoluble expression of TcB-TcC (Figure 6f). The same is true for the constriction site and the D-ring, where mutations of the highly conserved D34 and D505 residues to alanine or arginine do not cause a decrease in toxicity (Figure 6f, Supplementary Figure 9). These results indicate that single point mutations of conserved residues in the narrow passage of TcB-TcC are not sufficient to change the property of the respective region such that the translocation of the ADP-ribosyltransferase is negatively influenced. A combination of mutations is probably necessary to reach this. Unfortunately, insertion of several mutations at the same time seem to have a strong effect on the overall fold of the protein since it results in almost all cases in insoluble expression of TcB-TcC.

In summary, the structure of the C-terminal region of the ADP-ribosyltransferase in the narrow passage of TcB provides a snapshot of the translocation process, in which a temporary stabilization of the protein is achieved by transient interactions with residues of the channel wall.

### Pore formation of TcA in different lipid environments

The previously obtained cryo-EM structure and MD simulations of TcA in lipid nanodiscs showed that lipid headgroups intercalate between the TcA protomers (Gatsogiannis et al., 2016). The natural flexibility of this region resulted in a lower resolution not only in the structure of the TcA pore (Gatsogiannis et al., 2016), but also in the pore structures of ABC(D651A) and ABC(WT) (Supplementary Figure 2e, Supplementary Figure 5e).

To find out if membrane insertion of TcA is specific to certain lipids, we induced pore formation and checked how well the protein reconstituted into liposomes using different lipids. TcA integrated readily into 1-palmitoyl-2-oleoyl-phosphatidylcholine (POPC), di-oleoyl-phosphatidylcholine (DOPC), POPC with 20% cholesterol and brain polar lipids (BPL). In all cases, more than 70% of TcA is reconstituted in liposomes. The reconstitution efficiency in liposomes composed of POPC with 20% phosphatidylinositol (PI) or POPC with 20% sulfatidated lipids was slightly worse and when POPC with 20% 1-palmitoyl-2-oleoyl-phosphatidylglycerol (POPG) or a *E. coli* polar lipid extract (ECL) was used, considerably less proteoliposomes were formed (only 35-50% of the protein, Figure 7a,b). This demonstrates that although TcA has no pronounced affinity for certain lipids, it preferably integrates into membranes that do not contain negatively charged head groups. Furthermore, mammalian lipid mixtures seem to be preferred over bacterial lipids.

**Figure 7:**
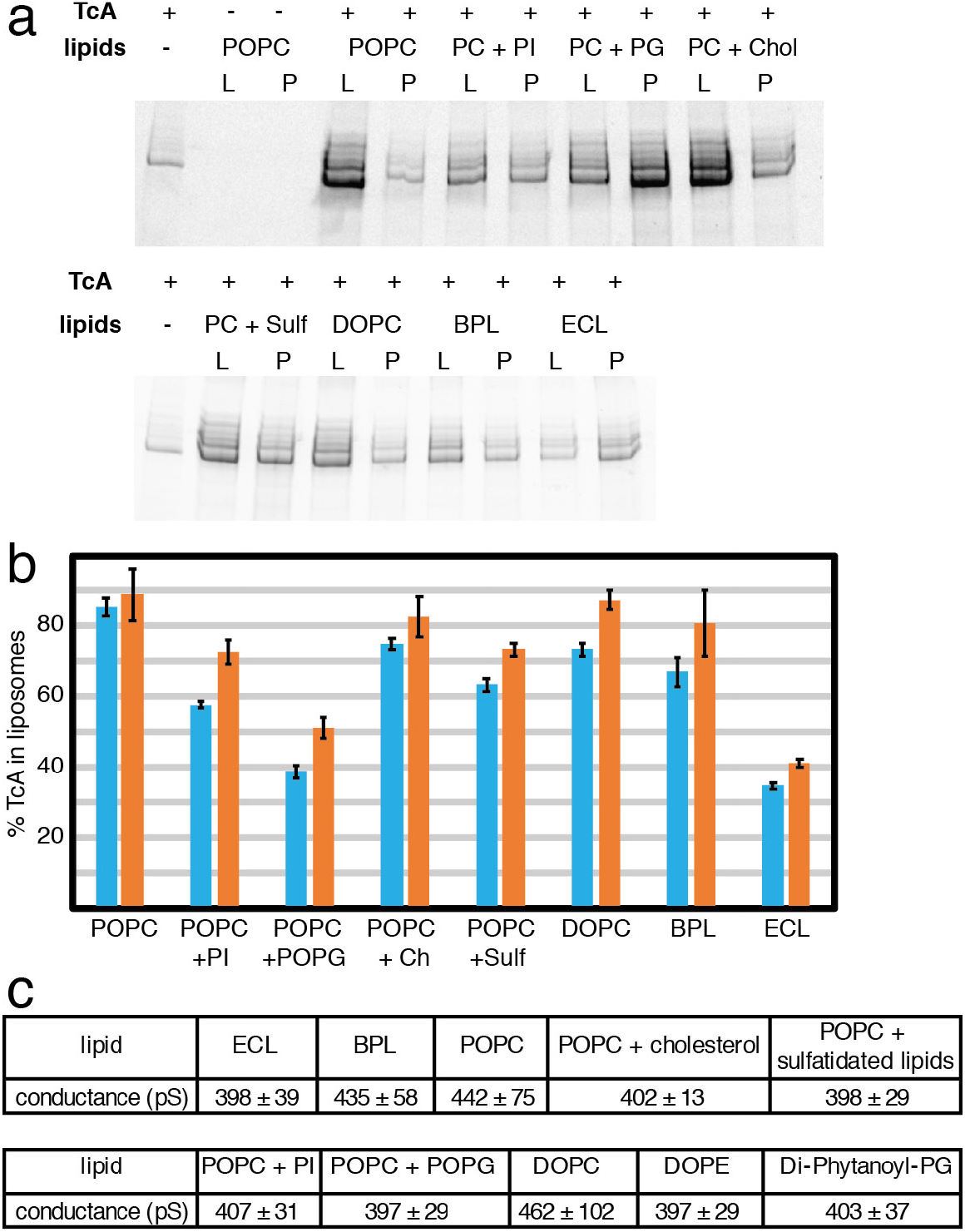
Lipid interaction and pore formation of TcA. **a**: Integration of TcA in liposomes of various compositions. SDS-PAGE of the proteoliposome flotation of TcA incubated with various lipids (POPC, POPC + 20% liver PI, POPC + 20% POPG, POPC + 20% cholesterol, POPC + 20% sulfatidated lipids, DOPC, brain polar lipid extract, *E. coli* polar lipid extract) at pH 11 for 48 h. L: liposome fraction, P: pellet fraction. **b**: Percentage of TcA in the liposome fraction in two independent liposome flotation experiments determined by densitometry of SDS-gel bands (blue and orange bars, respectively). The error bars represent standard deviations of three samples. **c**: Averages and standard deviations of single channel conductance values of TcA in different lipids in black lipid bilayer experiments (see Supplementary Figure 10).

To understand whether TcA forms indeed a pore in the different types of lipids used, we reconstituted TcA into black lipid bilayers composed of different lipids and measured the conductance of the channel. TcA readily integrated in all lipid bilayers, however, in contrast to the previously measured conductance of 500 – 600 pS for the open pore in diphytanoyl-phosphatidylcholine (Gatsogiannis et al., 2013), the mean conductance values were between 397 and 462 pS in all conditions tested, even with bilayers formed exclusively by lipids with negatively charged headgroups (diphytanoyl-phosphatidylglycerol) (Figure 7c, Supplementary Figure 10). In addition, in bilayers consisting of phosphatidylcholine only or BPL, there is a considerable fraction of larger pores with a conductance of up to 600 pS (Supplementary Figure 10b,c,h). While this indicates that TcA can indeed form pores in different lipid environments, it demonstrates that the lipids have an influence on the diameter of the TcA pore.

### An alternative order of holotoxin assembly

Since we have learned from the holotoxin pore structure that the TcB-TcC cocoon can easily be accommodated inside the basin formed by the shell domains of TcA, we asked ourselves whether TcB-TcC can also bind to TcA after it is already inserted into the membrane. To confirm this possibility, we first formed TcA pores in nanodiscs and then added TcB-TcC. We indeed obtained holotoxin pores, clearly demonstrating that the complex can also be assembled in this order (Figure 8a).

**Figure 8:**
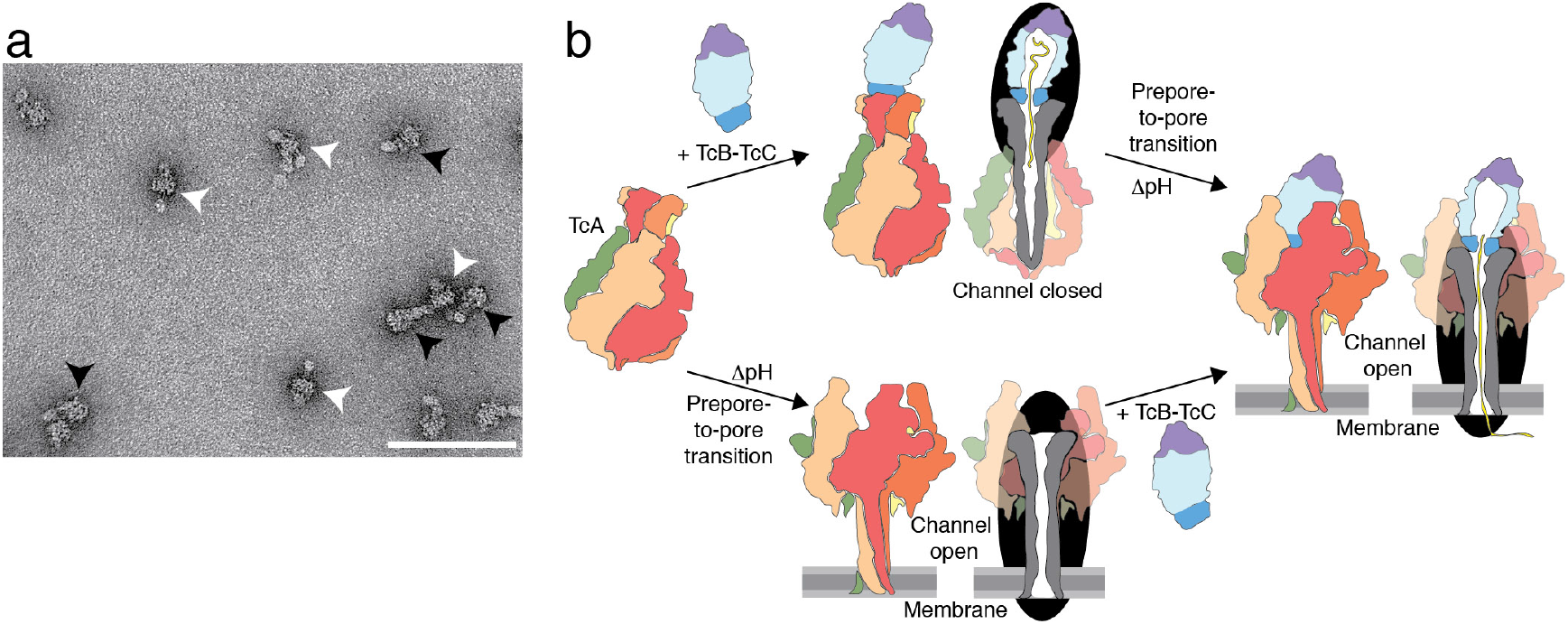
Model of two pathways of formation of the holotoxin pore. **a**: Negative stain electron micrograph of Tc holotoxin in nanodiscs formed by mixing TcA after the prepore-to-pore transition at pH 11 with TcB-TcC (80 and 500 nM, respectively), followed by size exclusion chromatography. Holotoxin prepores and pores are indicated with black and white arrowheads, respectively. Scale bar, 100 nm. **b**: Cartoon representation of Tc holotoxin assembly, target cell interaction and pore formation. The first pathway shows holotoxin formation with sub-nanomolar affinity from pentameric TcA and the TcB-TcC fusion protein (Gatsogiannis et al., 2018), followed by membrane association and pH-induced prepore-to-pore transition. The second pathway shows membrane association and prepore-to-pore transition of TcA, followed by holotoxin formation of the TcA pore and TcB-TcC. Subsequently, the toxic enzyme is translocated to the target cells with its C-terminus first.

Where and when Tc holotoxins assemble is still a matter of debate. The sub-nanomolar affinity of TcA and TcB-TcC (Gatsogiannis et al., 2018) suggests that the components are likely secreted as a fully assembled holotoxin. This is also supported by the finding that TcC is needed for the proper secretion of TcA and TcB from *P. luminescens* (Yang and Waterfield, 2013). However, what stands against the secretion as a fully assembled complex is the relatively large size of the holotoxin (1.7 MDa). Our results demonstrate that alternative routes of holotoxin complex formation are possible and happen likely *in vivo* (Figure 8b).

## Conclusion

The pore structure of the Tc holotoxin complex from *P. luminescens* at near-atomic resolution presented in this study allowed us to clarify mechanistic details of the prepore-to-pore transition of the assembled complex and toxin translocation. During pore formation of ABC, the TcB-TcC cocoon is pulled into a basin formed by the outer shell of the TcA protomers, which slightly adapt to fit the cocoon. This adjustment causes a break in the symmetry of TcA, but no structural change at the interface between TcB and TcA (Supplementary Movie 6). The structural flexibility of the upper part of the outer shell of TcA allows the ABC holotoxin to be assembled after TcA has already entered the membrane (Figure 8).

In cleavage-deficient ABC(D651A), the ADP-ribosyltransferase is still present inside the holotoxin, indicating that its full release from the cocoon requires autoproteolytic cleavage. Its major part remains unstructured in the TcB-TcC cocoon as in our previous structures of the holotoxin in its prepore state (Gatsogiannis et al., 2018; Meusch et al., 2014). However, the improved resolution of the present structure allowed us to model two regions of the ADP-ribosyltransferase for the first time. The first region is the N-terminus, which follows the aspartyl autoprotease site. It forms a loop that keeps the cleavage site between the catalytic aspartates. The second part is a 13 residue-long stretch close to the C-terminus that resides in the narrow passage of TcB below the constriction site. The C-terminal stretch is in an unfolded conformation and interacts with alternating charged and hydrophobic regions of the channel. A similar pattern has been described for chaperones (Morán Luengo et al., 2018) and the translocation pore of anthrax protective antigen (PA) (Jiang et al., 2015) (Supplementary Figure 11). There, the so-called ϕ-clamp forms a narrow (∼6 Å) hydrophobic constriction site (Jiang et al., 2015). How the substrates interact with the ϕ-clamp during translocation is not known in detail. It has been proposed that it seals the translocation pore around the translocated polypeptide and helps to protect hydrophobic patches in the translocated protein (Jiang et al., 2015; Krantz et al., 2005). The same is probably true for the Tc complex, however, a seal is not needed in this case since in contrast to PA, the Tc translocation channel is closed by the TcB-TcC cocoon on one side. Our results demonstrate that translocation happens spontaneously *in vitro*. It does not require driving forces such as pH gradients or chaperones, although folding of the toxin inside the host cell cytoplasm *in vivo* is likely supported by the chaperones of the host cell (Lang et al., 2014) (Ernst et al., 2017), and translocation might be accelerated by a pH gradient. Based on our previous results (Gatsogiannis et al., 2016), we have speculated that the direct entanglement with lipid head groups at the conserved tip of the TcA channel inside the membrane is partly responsible for the host specificity of Tc toxins. However, the results presented here demonstrate that this is likely not the case. While the type of lipid has an influence on the diameter of the TcA pore, it does not interfere with pore formation in general.

## Acknowledgements

We thank A. Elsner and K. Vogel-Bachmayr for purification of TcdA1 and TcdB2-TccC3, respectively. We acknowledge the help of D. Pan and F. Müller in preparing the cross-linked TcB-TcC(WT) complex. We thank A. Brockmeyer and M. Metz for performing mass spectrometry and the annotation of the cross-linked peptides. This work was supported by funds from the Max Planck Society (to S.R.) and the European Research Council under the European Union’s Seventh Framework Programme (FP7/2007-2013) (grant no. 615984) (to S.R.).

## Author Contributions

S.R. designed and supervised the project. D.R. prepared the holotoxin in nanodiscs, prepared cryo-EM samples, processed and analyzed cryo-EM data and performed all biochemical experiments. O.H. recorded the cryo-EM images. R.B. and D.R. performed single-channel conductivity experiments. D.R. prepared figures, D.R. and S.R. wrote the manuscript.

## Author Information

The cryo-EM densities of ABC(D651A) and ABC(WT) pore states have been deposited in the Electron Microscopy Data Bank under accession numbers … and …, respectively. Coordinates of ABC(D651A) and ABC(WT) pore states have been deposited in the Protein Data Bank under accession number…and …, respectively. Correspondence and requests for materials should be addressed to S.R.

## Supplementary Figures

**Supplementary Figure 1:**
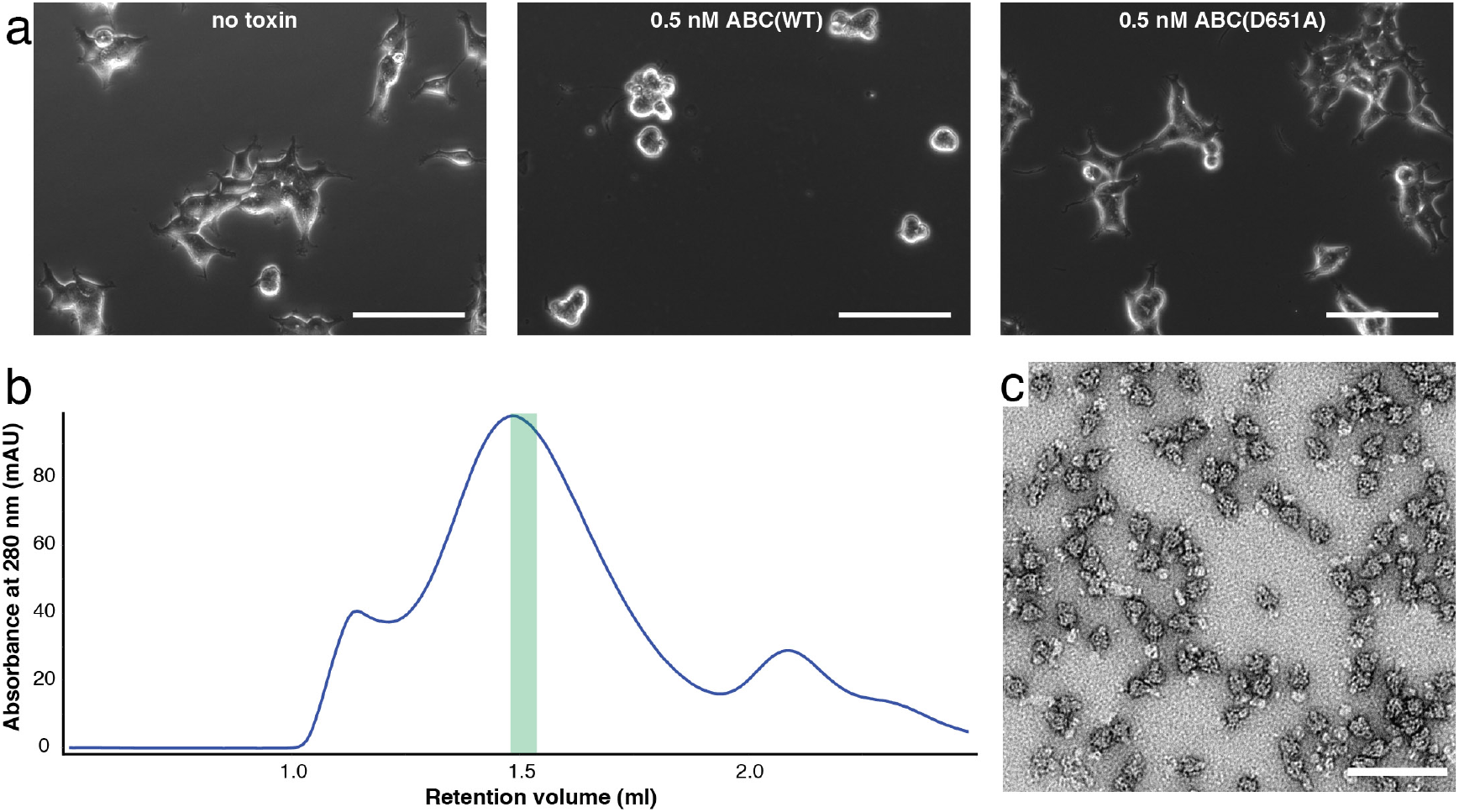
Preparation of ABC(D651A) in nanodiscs. **a**: Intoxication of HEK 293T cells with ABC(WT) and ABC(D651A). 2×10^4^ cells in DMEM/F12 medium were incubated with PBS or 0.5 nM of ABC for 16 h at 37 °C before imaging. Experiments were performed in triplicates with qualitatively identical results. Scale bars, 100 µm. **b**: Size exclusion chromatography of ABC(D651A) after integration in MSP5ΔH5-POPC nanodiscs. The main peak fraction indicated in green was used for cryo-EM. **c**: Negative-stain electron micrograph of the peak fraction indicated in **a**, showing predominantly holotoxin pores. Scale bar, 100 nm.

**Supplementary Figure 2:**
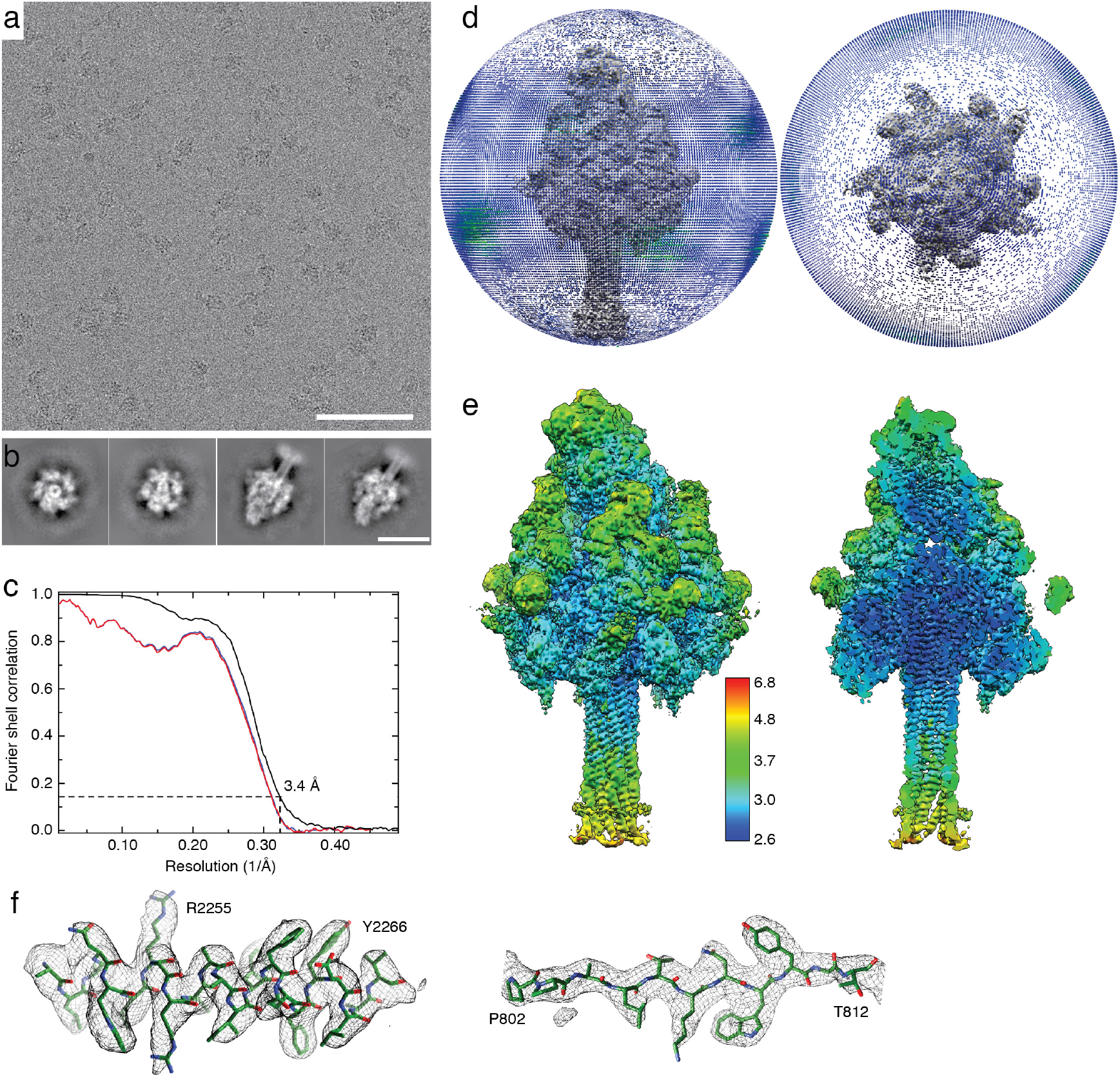
Cryo-EM of ABC(D651A) in a nanodisc. **a**: Representative electron micrograph of vitrified ABC(D651A) at 2.1 µm defocus and 100 e^−^/Å^2^ total dose acquired with a Falcon 3 direct electron detector. Scale bar, 100 nm. **b**: Representative 2D class averages. Top views (left two images) and side views (right two images) are shown. Scale bar, 20 nm. **c**: Fourier shell correlation (FSC) of the obtained cryo-EM map (black curve) and map-to-model FSCs of two independently refined half maps (red and blue curves, respectively). The dashed line shows the 0.143 FSC cutoff criterion. **d**: Angular distribution plots of the final round of refinement in side view and top view. Each stick represents a projection view. The size and color of sticks (blue-to-green) is proportional to the number of particles. **e**: Local resolution estimation of the cryo-EM density map, side view (left) and section (right), colored according to local resolution. **f**: Superimpositions of representative areas of the cryo-EM density maps and the models, shown for an α-helical part in TcA (left) and a β-strand in TcB (right).

**Supplementary Figure 3:**
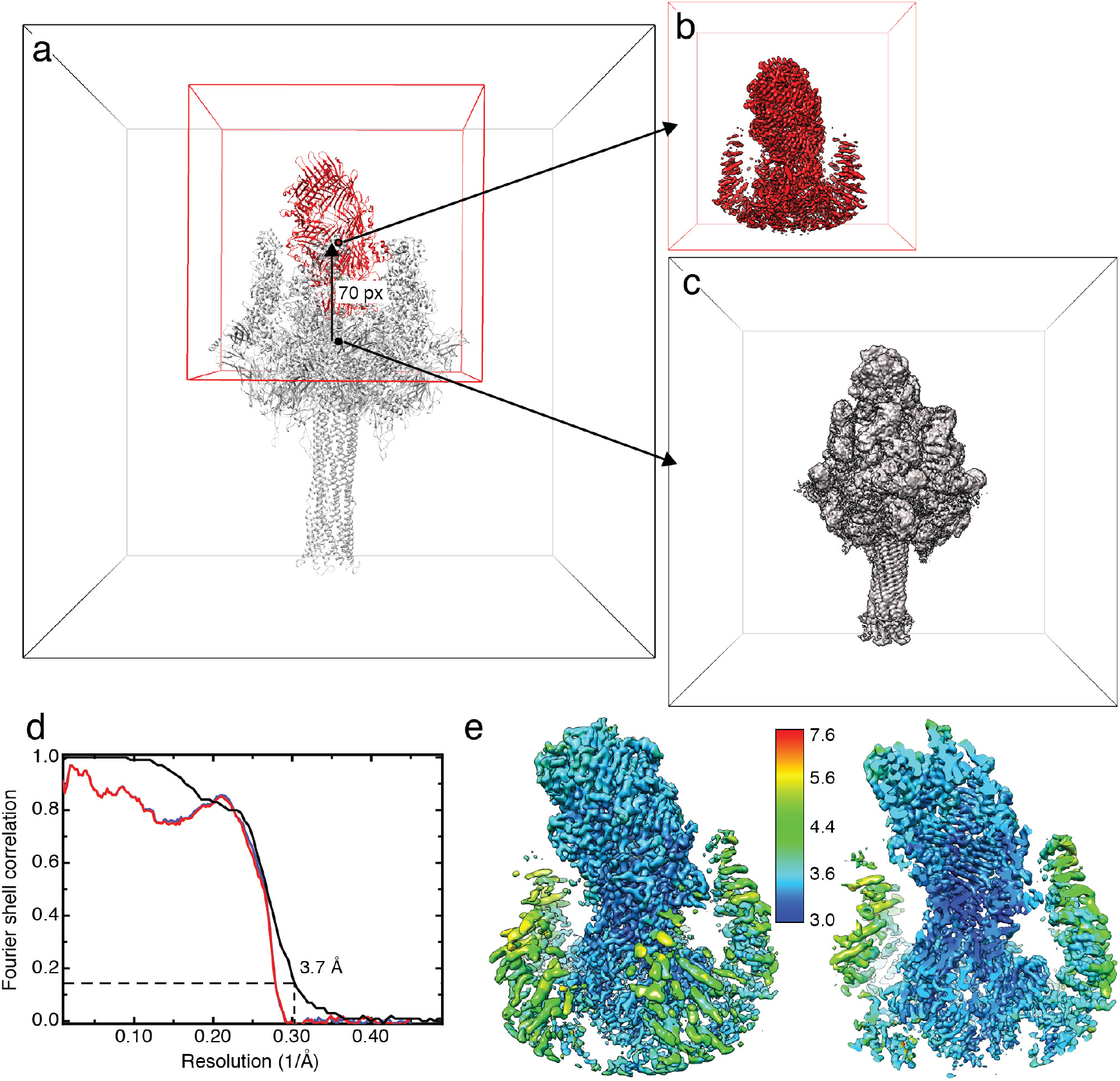
Shift of the particle box center from the center of ABC(D651A) to the center of TcB-TcC. **a**: Scheme of shift of box center from the center of the ABC(D651A) holotoxin (black dot) to the center of TcB-TcC (red dot) by 70 pixels (corresponds to 78 Å). The box was re-scaled from 384 pixels to 200 pixels. The model of the holotoxin is shown in the box for comparison, with TcA in gray and TcB-TcC in red. **b,c**: Cryo-EM maps of the final refinement of the shifted reconstruction (**b**, red) and the original reconstruction of the full ABC(D651A) holotoxin (**c**, gray). The particle boxes of the maps are also depicted. **d**: FSC of the cryo-EM map (black curve) and map-to-model FSCs of the model of TcB-TcC with two independently refined half maps (red and blue curves, respectively). The dashed line shows the 0.143 FSC cutoff criterion. **e**: Local resolution estimation of the cryo-EM map, side view (left) and section (right), colored according to local resolution.

**Supplementary Figure 4:**
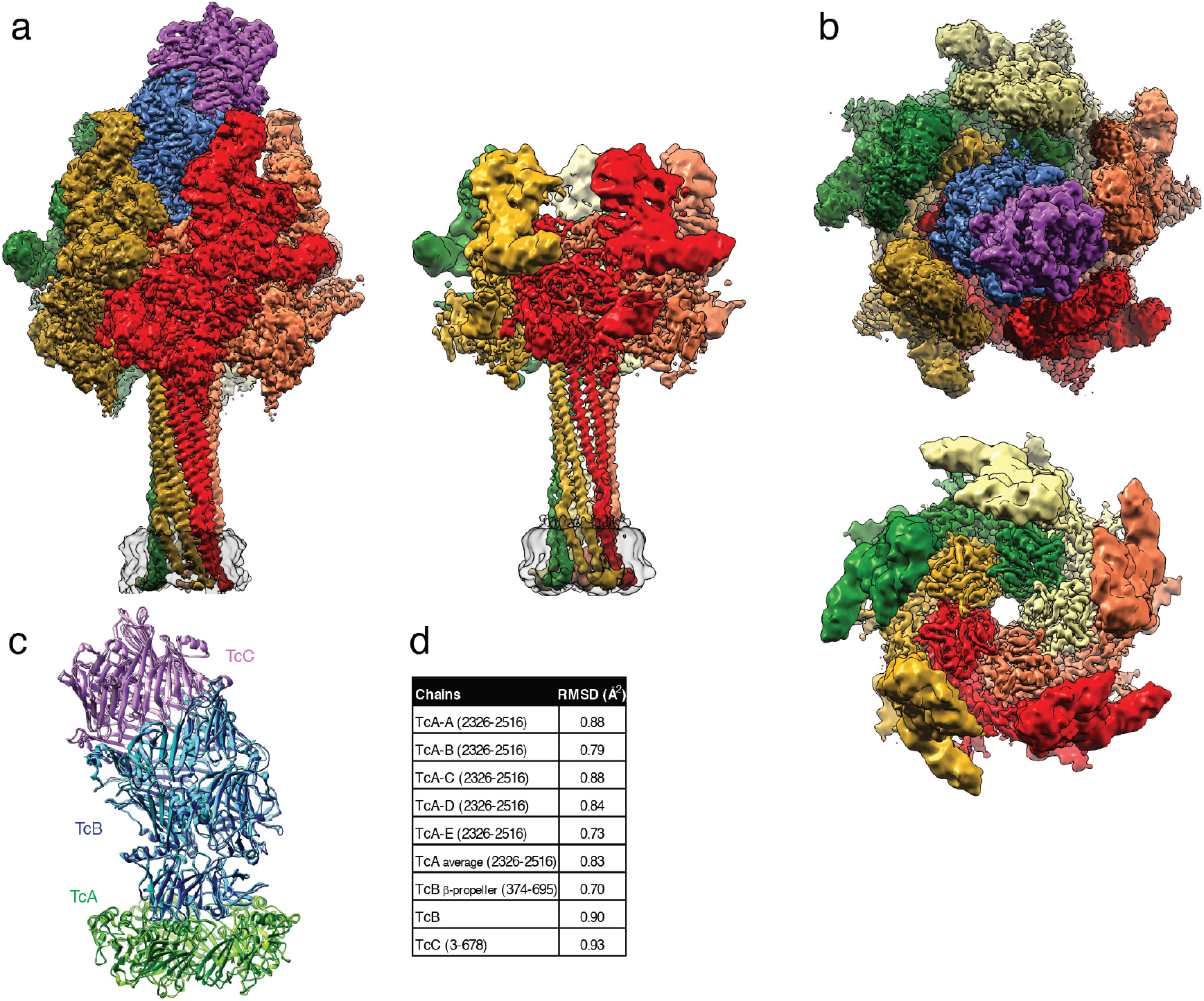
Comparison of cryo-EM maps of ABC(D651A) and TcA. **a**: Side view of the density maps of ABC(D651A) and TcA (EMDB 4068) embedded in lipid nanodiscs. **b**: Top views of the maps in panel **a**. The individual subunits and the nanodiscs are colored like in Figure 1a. **c**: Overlay of models of the TcB-TcC and TcA (residues 2327-2516) of the prepore of ABC(D651A) (pdb ID 6H6F, light colors) with the respective model of the pore of ABC(D651A) (dark colors). The TcB-binding domains of TcA were aligned on each other. **d**: Backbone RMSDs of the individual TcA chains, all TcA chains combined, the β-propeller domain of TcB, TcB and TcC between the prepore and pore form of ABC(D651A). RMSD values were calculated in Chimera.

**Supplementary Figure 5:**
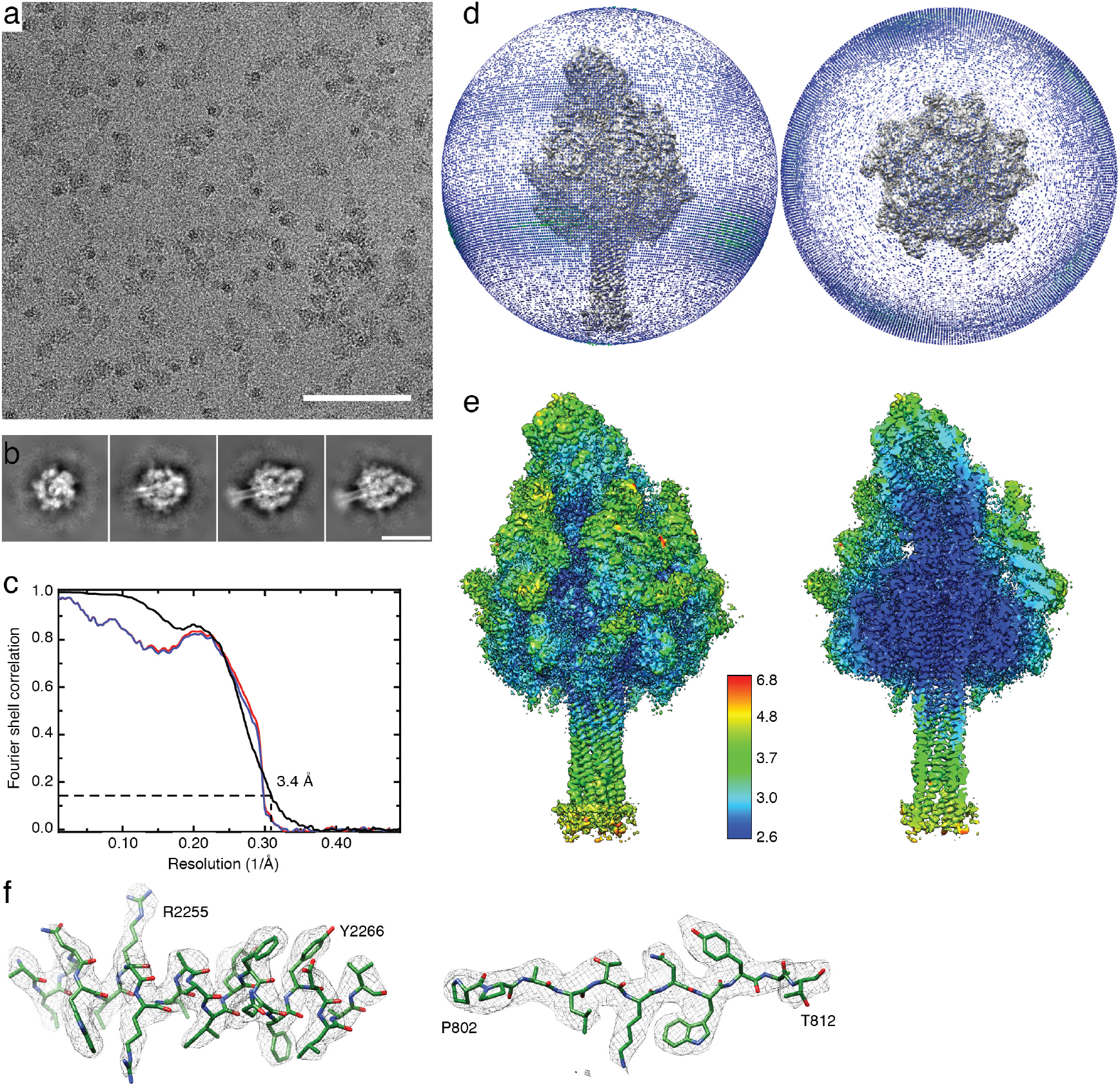
Cryo-EM of ABC(WT) in a nanodisc. **a**: Representative electron micrograph of vitrified ABC(WT) at 2.0 µm defocus and 63 e^−^/Å^2^ total dose acquired with a K2 direct electron detector. Scale bar, 100 nm. **b**: Representative 2D class averages. Top and tilted views (left two images) and side views (right two images) are shown. Scale bar, 20 nm. **c**: Fourier shell correlation (FSC) of the obtained cryo-EM map (black curve) and map-to-model FSCs of two independently refined half maps (red and blue curves, respectively). The dashed line shows the 0.143 FSC cutoff criterion. **d**: Angular distribution plots of the final round of refinement in side view and top view. Each stick represents a projection view. The size and color of sticks (blue-to-green) is proportional to the number of particles. **e**: Local resolution estimation of the cryo-EM map, side view (left) and section (right), colored according to local resolution. **f**: Superimpositions of representative areas of the cryo-EM density maps and the models, shown for an α-helical part in TcA (left) and a β-strand in TcB (right).

**Supplementary Figure 6:**
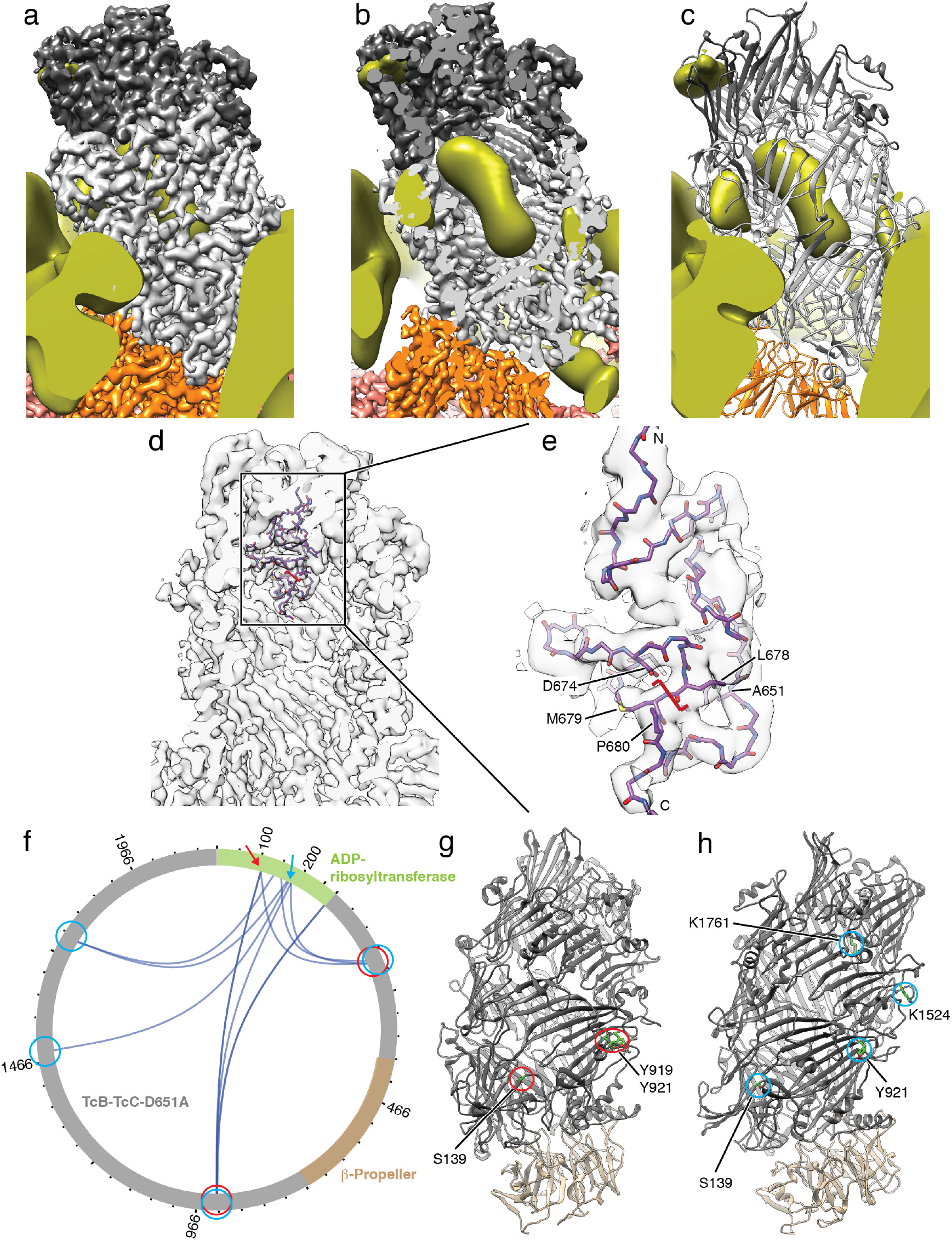
Variability analysis of the TcB-TcC cocoon, visualization of the aspartyl protease domain of TcC and XL-MS analysis of ABC(D651A) with encapsulated, non-cleaved toxic enzyme. **a-c**: 3D variability analysis of the ABC(D651A) structure. The 3D average and variance map were created in SPHIRE (Moriya et al., 2017) with the input images filtered to 25 Å. The 3D variance map is shown in gold. a,b: Overlay of the cryo-EM density map of ABC(D651A) (colored in pale red, orange, light gray and dark gray according to TcA, TcB β-propeller, TcB cocoon and TcC, respectively) and 3D variance map in side view (**a**) and section (**b**). **c**: Model of TcB-TcC with the 3D variance map (gold). Note the highest variance in the center of the cocoon and at the upper domains of TcA. **d**: Position of the aspartyl autoprotease domain (residues 618 – 678 of TcC) with the attached N-terminus of the toxic enzyme (residues 679 – 683 of TcC) in the TcB-TcC cocoon. The model is overlaid with the cryo-EM map obtained by shifting the center of reconstruction (Supplementary Figure 3). The cleavage site is indicated (red bracket). **e**: Detailed view of panel **d**. The protein is cleaved between L678 and M679. The residues forming the aspartyl autoprotease cleavage site (D674 and A651, which is D651 in WT) and the conserved P680 are indicated. **f**: Visualization of crosslinks (dark blue curves) in TcB-TcC-D651A between the TcB-TcC cocoon (gray, β-propeller in sand) and the ADP-ribosyltransferase (green). Two individual positions (K107 and K175, K178, K185) and their crosslinks to the TcB-TcC cocoon are indicated with red and blue arrows and circles, respectively. All crosslinks with a score of 50 and higher are shown. The plot was created using xVis (Grimm et al., 2015). A complete list of all crosslinks is shown in Supplementary Table 3. **g,h**: The TcB-TcC cocoon with all amino acids that crosslink with K107 indicated and highlighted with red circles (**g**) and crosslinks with K175, K178, K185 indicated and highlighted with blue circles (**h**). Residue numbering according to pbd ID 6H6F.

**Supplementary Figure 7:**
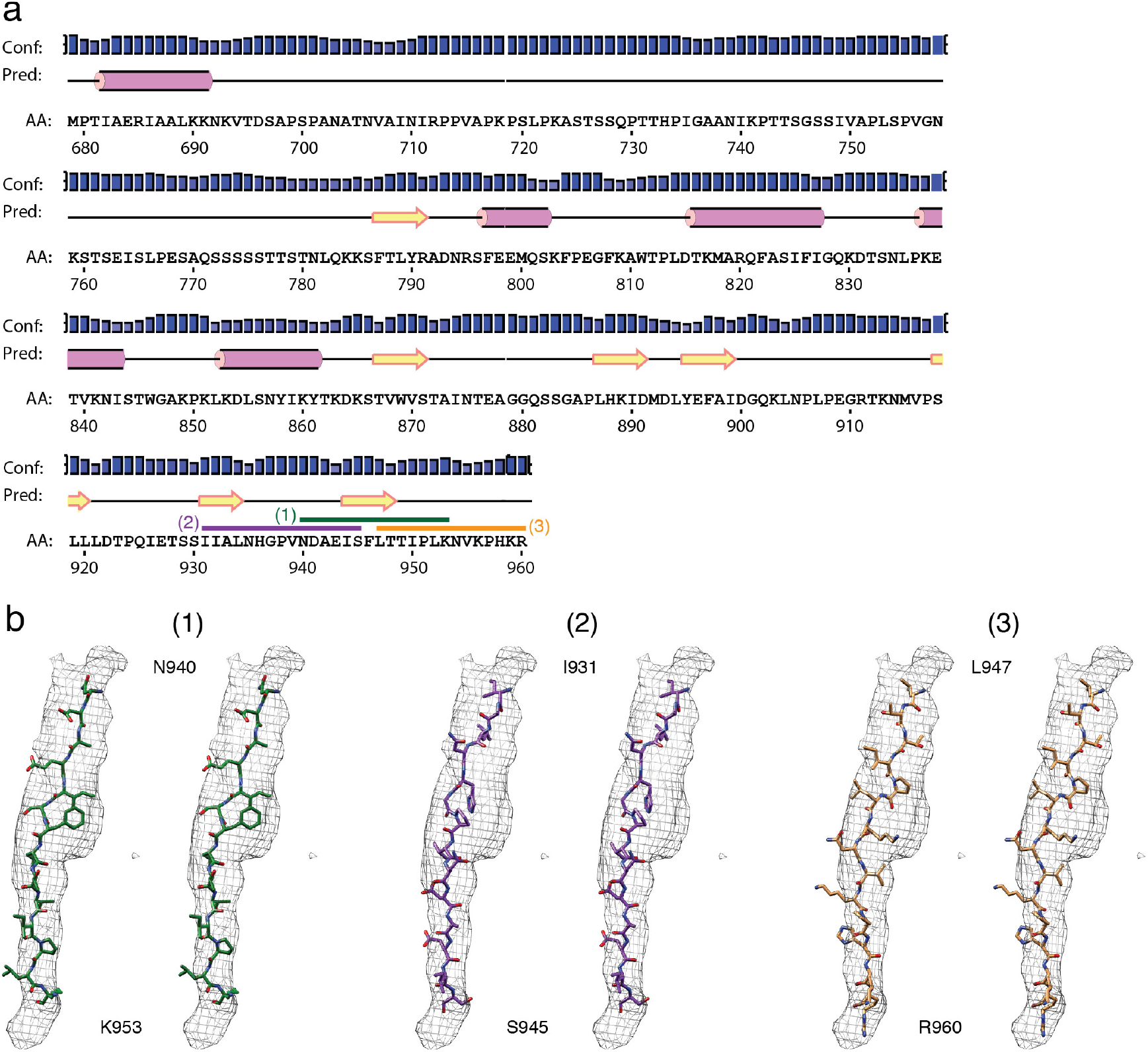
Comparison of different models close to the C-terminus of the toxic enzyme in the density in the narrow passage of TcB. **a**: Secondary structure prediction of the ADP-ribosyltransferase of TccC3, obtained by PSIPRED (Buchan et al., 2013). Predicted α-helices are shown as purple cylinders, β-strands are shown as yellow arrows. The positions of modeled peptides in b are shown as green, purple and orange bars. Residue numbering starts with the first residue after the cleavage site of TcC (M679). Conf: confidence, Pred: prediction, AA: residue. **b**: Stereo views of three atomic models of possible peptides close to the C-terminus of the ADP-ribosyltransferase in the cryo-EM density (shown as mesh representation) in the narrow passage of TcB. N940 – K953 (green) fits best, with the bulky density in the center occupied by F946.

**Supplementary Figure 8:**
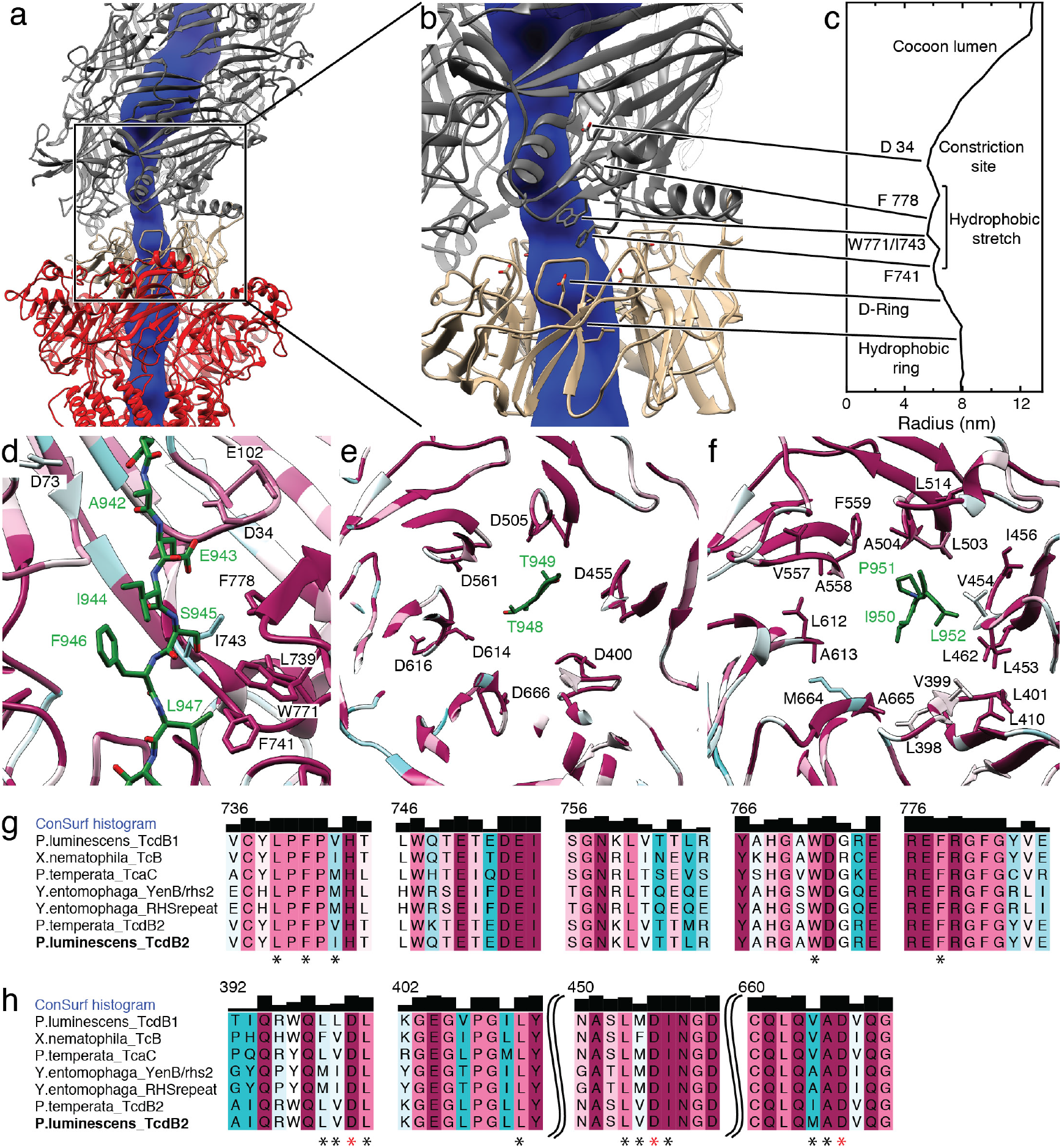
Analysis of translocation channel of TcB and conservation plot of residues. **a**: The central translocation channel of TcB, spanning from the cocoon (gray) through the TcB β-propeller (sand) to TcA (red). The channel lumen obtained by ChExVis (Masood et al., 2015) is shown in blue. **b**: Section of panel **a**, showing constrictions in the TcB β-propeller and cocoon. D34 in the constriction site and residues of the hydrophobic stretch, the D-ring and the hydrophobic ring (see Figure 6) are shown. **c**: Plot of inner radius of the TcB channel. Residues and motifs forming constrictions are indicated. d-f: The constriction site and the hydrophobic stretch (**d**), the negatively charged ring (**e**) and the hydrophobic ring (**f**) with mapped sequence conservation ranging from minimum (cyan) to maximum (magenta). The translocating ADP-ribosyltransferase is depicted in green. Views and highlighted residues are the same like in Figure 6c-e. **g**: Sequence alignment of the hydrophobic stretch of TcB. Residues shown in panel d are indicated (*). **h**: Sequence alignment of selected regions of the D-ring and the hydrophobic ring. Residues shown in panels e and f are indicated by red and black asterisks, respectively.

**Supplementary Figure 9:**
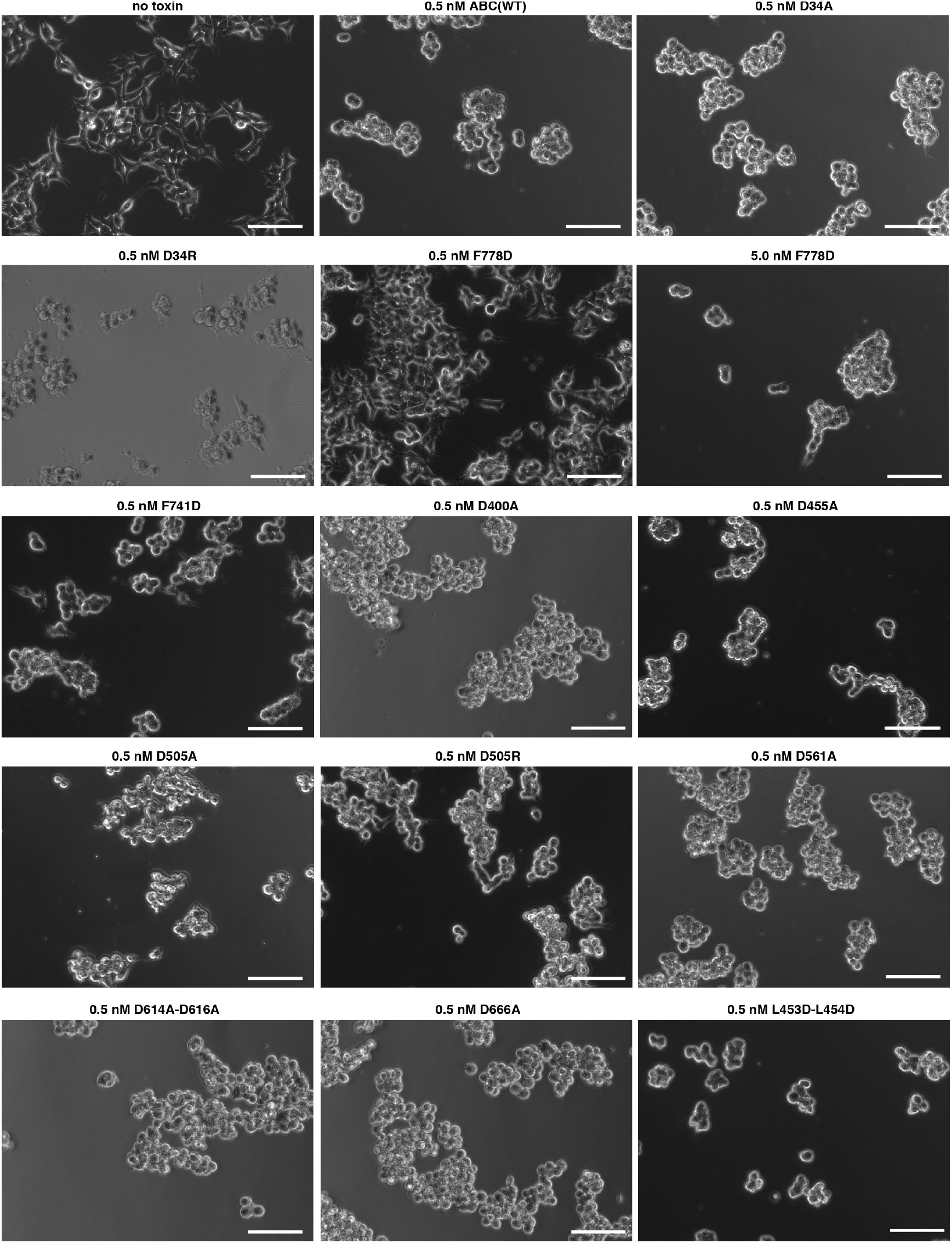
Comparison of toxicity of TcB variants. Intoxication of HEK 293T cells with holotoxins formed by TcA(WT) and the indicated TcB-TcC variants. Intoxicated cells round up and detach from the surface. 2×10^4^ cells in DMEM/F12 medium were incubated with 0.5 nM or 5.0 nM of ABC for 16 h at 37 °C before imaging. Experiments were performed in duplicates with qualitatively identical results. Scale bars, 100 μm.

**Supplementary Figure 10:**
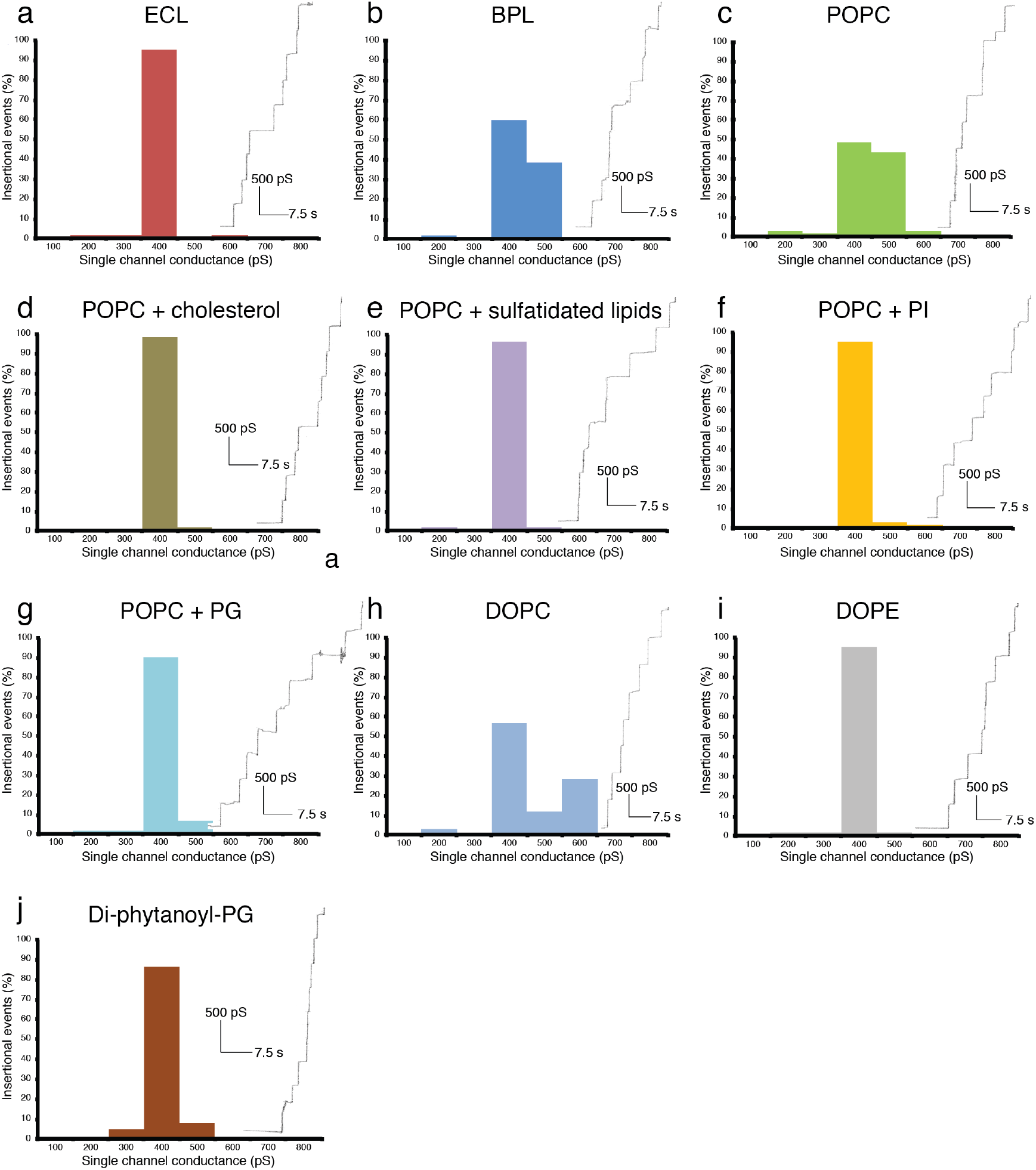
Single channel conductance of TcA in different lipids. All histograms were constructed from the data of 60 pore insertional events for each condition. For each experiment, an exemplary recording of the single-channel current versus time is shown next to the histogram. **a**: TcA and *E. coli* polar lipid extract (ECL). **b**: TcA and brain polar lipid extract (BPL). **c**: TcA and POPC. **d**: TcA and POPC with 20% cholesterol. e: TcA and POPC with 20% sulfatidated lipids. **f**: TcA and POPC with 20% liver phosphatidylinositol (PI). **g**: TcA and POPC with 20% phosphatidylglycerol (PG). **h**: TcA and di-oleoyl-phosphatidylcholine (DOPC). **i**: TcA and di-oleoyl-phosphatidylethanolamine (DOPE). **j**: TcA and di-phytanoyl-phosphatidylglycerol.

**Supplementary Figure 11:**
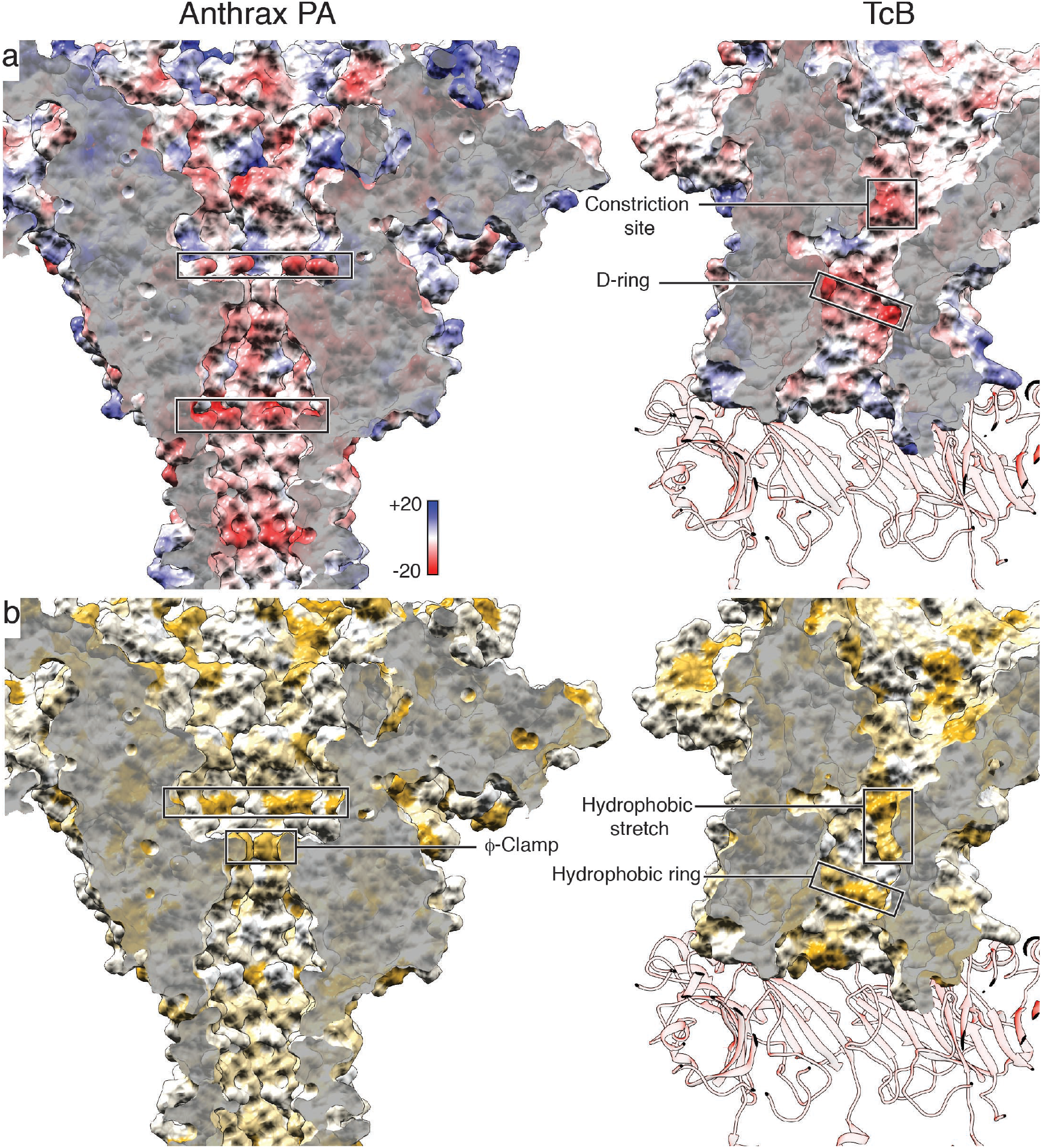
Comparison of the Coulomb potentials and surface hydrophobicities of the translocation channels of anthrax protective antigen (PA) with the narrow passage of TcB. **a**: Surface representation of anthrax PA (left) and TcB (right), colored according to the Coulomb potential (kcal mol^−1^ e^−1^) at pH 7.0. The interacting parts of TcA at the bottom are shown as transparent ribbon representation. The negatively charged regions of both structures close to the channel constrictions are highlighted in boxes. Translocation direction is from top to bottom. **b**: The same view as in panel a, colored according to hydrophobicity. Hydrophobic regions are colored ochre, non-hydrophobic regions are colored white. Hydrophobic regions inside the channels are highlighted in boxes.

## Movie legends

Supplementary Movie 1: Cryo-EM density map of the ABC(D651A) pore with the fitted atomic model. Representative areas of the cryo-EM density in TcA and TcB-TcC are shown.

Supplementary Movie 2: Morph between TcA-A (red) and the other four TcA chains, visualizing the structural changes in the upper part of the α-helical shell in comparison to the rest of the molecule.

Supplementary Movie 3: Comparison between the ABC and ABC(D651A) pore by moving through vertical slices along the central axis of the molecules. TcA, TcB and TcC are shown as ribbon representations. The nanodisc is shown in transparent gray. Additional density not corresponding to the atomic models is filtered to 10 Å and shown in dark gray.

Supplementary Movie 4: Cryo-EM density map of the ABC(D651A) pore after shifting the center of reconstruction. The atomic model of TcB-TcC (blue and purple, respectively) is fitted into the map. In the second part, the movie zooms in and focuses on the aspartyl protease domain of TcC.

Supplementary Movie 5: Cryo-EM density map inside the narrow passage of TcB with the different C-terminal polypeptides of the ADP-ribosyltransferase fitted into the density (Supplementary Figure 7b).

Supplementary Movie 6: Morph of the cryo-EM density of the ABC(D651A) prepore (EMDB 0150) to the density of the ABC(D651A) pore in a nanodisc. The morph between the densities is followed by a morph between the states using the atomic models of the proteins, showing the prepore-to-pore transition on the level of the holotoxin and the opening of the translocation channel in the membrane. The outer shell, the linker and the transmembrane channel of TcA are shown in red, black and yellow, respectively. TcB is shown in blue, and TcC is shown in purple.

## Methods

### Protein production

*P. luminescens* TcdA1 (TcA) was expressed in *E. coli* BL21-CodonPlus (DE3)-RIPL in LB medium and purified as described previously (Gatsogiannis et al., 2013). Fusion proteins *P. luminescens* TcdB2-TccC3 (TcB-TcC(WT)) and cleavage-deficient TcdB2-TccC3 variant D651A (TcB-TcC(D651A)) were also expressed in *E. coli* BL21-CodonPlus (DE3)-RIPL in LB medium and purified as described previously (Gatsogiannis et al., 2016), as well as all TcB-TcC mutants. The nanodisc scaffold protein MSP1D1 was expressed and purified using pMSP1D1 (Addgene #20061) as described previously (Denisov et al., 2004).

### Holotoxin formation and nanodisc integration

TcdA1 and TcdB2-TccC3(WT) or TcdB2-TccC3(D651A) were mixed with a 1.5-fold excess of TcB-TcC over TcA pentamer and subsequently the excess of TcB-TcC was separated from the holotoxin by size exclusion chromatography (SEC) on a Superose 6 5/150 column (GE Life Science). For ABC(D651A), the holotoxin was concentrated to 1 mg/ml and mixed with a 12-fold molar excess of pre-formed MSP5ΔH5-POPC nanodiscs (Cube Biotech) in 20 mM Tris-HCl pH 8.0, 250 mM NaCl, followed by 1 h of incubation on ice. For ABC(WT), nanodiscs were pre-formed using MSP1D1 and POPC in a 1:55 molar ratio. The holotoxin was concentrated to 1.2 mg/ml and mixed with a 10-fold molar excess of nanodiscs in 20 mM Tris-HCl pH 8.0, 250 mM NaCl, followed by 1 h of incubation on ice. Subsequently, the holotoxin/nanodisc mixtures were dialyzed against 20 mM CAPS-NaOH pH 11.2, 250 mM NaCl for 2 days at 4 °C. After dialysis, the holotoxin/nanodisc mixtures were subjected to SEC on a Superose 6 5/150 column equilibrated in 20 mM Tris-HCl pH 8.0, 250 mM NaCl, and the main peak containing ABC in nanodiscs was used for subsequent cryo-EM (Supplementary Figure 1b,c).

### Liposome overlay assay

TcA (1 mg/ml) was mixed with different lipid mixes dissolved in 5% β-D-octylglucoside (Anatrace) with a total lipid concentration of 5 mg/ml. We tested POPC, POPC + 20% liver PI, POPC + 20% POPG, POPC + 20% cholesterol, POPC + 20% sulfatidated lipids (porcine brain sulfatides), DOPC, BPL and ECL. All lipids were purchased from Avanti Polar Lipids (Alabaster, AL 35007). After 1 h of incubation of TcA and lipids dissolved in OG at 4 °C and pH 8, the mixtures were dialyzed against 20 mM CAPS-NaOH pH 11.2, 150 mM NaCl for 2 days at 4 °C. Subsequently, the dialyzed proteoliposomes were mixed 1:1 with 60% glucose in dialysis buffer to 100 µl total volume, overlaid with 125 μl 25% glucose and 30 μl dialysis buffer and centrifuged at 175,000×g for 4 h at 4 °C in a Beckman TLA 120.1 rotor. After centrifugation, the top and bottom fractions of the sucrose gradient were analyzed via SDS-PAGE and the respective TcA amounts were quantified via band densitometry.

### Single channel conductance measurements

We measured the single channel conductance of TcA with different lipid compositions with black lipid membrane experiments. All lipids were obtained from Avanti Polar Lipids. The instrumentational setup consisted of a Teflon chamber with two compartments of 5 ml each which are connected by a small hole with a surface area of 0.4 mm^2^. The membrane current was measured with a pair of Ag/AgCl electrodes with salt bridges. The electrodes were switched in series with a voltage source and homemade current amplifier on the basis of Burr Brown operational amplifier as described previously (Gatsogiannis et al., 2013). Solutions of different lipids and lipid mixes (Supplementary Figure 10) at 10 mg/ml total lipid concentration were prepared in 80% n-decane/20% butanol. The lipid solutions were “painted” across the hole, resulting in membrane formation. After the membrane turned black, 20 pM of TcA were added to the cis side (the black side) of the chamber. All measurements were performed with a membrane potential of 20 mV in 20 mM CAPS-NaOH pH 11, 1 M KCl.

### Sample vitrification and cryo-EM data acquisition

For ABC(D651A), 2.4 μl of 0.12 mg/ml of the protein preparation were applied to glow-discharged holey carbon grids (Quantifoil, QF 2/1, 300 mesh) covered with a 2 nm carbon layer. 1.2 μl of 0.1% Tween-20 was added to the sample on the grid and blotting was performed immediately afterwards using a Cryoplunge3 (Cp3, Gatan) with 2.2 s blotting time at 20 °C and 90% humidity. For ABC(WT), 2.8 µl of 0.10 mg/ml of the protein preparation were applied to glow-discharged holey carbon grids (Quantifoil, QF 2/1, 300 mesh) covered with a 2 nm carbon layer. Blotting was performed using a Vitrobot Mark IV (Thermo Fisher) with 2.5 s blotting time at 5 °C and 100% humidity.

For ABC(D651A), two data sets were collected at a Cs-corrected Titan Krios EM (Thermo Fisher) equipped with an XFEG and a Falcon III direct electron detector. Images were recorded using the automated acquisition program EPU (FEI) at a magnification of 59,000, corresponding to a pixel size of 1.11 Å/pixel on the specimen level. 12,040 movie-mode images were acquired in total in a defocus range of 1.2 to 2.6 μm. Each movie comprised 40 frames acquired over 1.5 s with a total cumulative dose of **∼** 100 e^−^/Å^2^. The data sets were merged for image processing.

For ABC(WT), one data set was collected at a Cs-corrected Titan Krios EM (Thermo Fisher) equipped with an XFEG and a K2 direct electron detector (Gatan). Images were recorded using the automated acquisition program EPU (FEI) at a magnification of 130,000, corresponding to a pixel size of 1.05 Å/pixel on the specimen level. 2,660 movie-mode images were acquired in a defocus range of 1.4 to 2.6 μm. Each movie comprised 40 frames acquired over 15 s with a total cumulative dose of **∼** 61 e^−^/Å^2^.

### Image processing

After initial screening, 8,903 integrated images were selected for further processing for ABC(D651A), and 2,053 integrated images were selected for further processing for ABC(WT). Movie frames were aligned, dose-corrected and averaged using MotionCor2 (Zheng et al., 2017). CTF parameters were estimated using CTER (Penczek et al., 2014), implemented in the SPHIRE software package (Moriya et al., 2017). Outlier images were removed using the graphical CTF assessment tool in SPHIRE (Moriya et al., 2017). For ABC(D651A), 2,400 particles were manually picked initially and 2D class averages used as an autopicking template were generated using ISAC (Yang et al., 2012) in SPHIRE. 619,594 particles were auto-picked from the images using Gautomatch (Zhang et al., 2011) and extracted with a box size of 384 pixels. For ABC(WT), 390,353 particles were auto-picked using crYOLO (Wagner et al., 2018) using a box size of 300 pixels. 269,142 particles were subsequently extracted using a box size of 420 pixels.

Reference-free 2D classification and cleaning of the dataset was performed with the iterative stable alignment and clustering approach ISAC in SPHIRE. ISAC was performed with a pixel size of 5.61 Å/pixel for ABC(D651A) and 5.80 Å/pixel for ABC(WT) on the particle level. The ‘Beautifier’ tool of SPHIRE was then applied to obtain refined and sharpened 2D class averages at the original pixel size, showing high-resolution features (Supplementary Figure 2b, 5b). Only particles in ISAC classes showing a full holotoxin in the pore form were kept for 3D refinement, resulting in 337,823 particles for ABC(D651A) and 64,806 particles for ABC(WT), respectively.

We used the cryo-EM density map of the TcdA1 pore (EMD-4068 (Gatsogiannis et al., 2016)), manually docked with a molecular map of TcB-TcC (pdb ID 4O9X) and filtered to 25 Å, as initial reference for 3D refinement. 3D refinement without imposing symmetry was performed in SPHIRE using MERIDIEN (Moriya et al., 2017), using 7.5° initial angular sampling. 3D classification using SORT3D in SPHIRE, based on the final projection parameters of MERIDIEN, resulted in classes with different orientations of the asymmetric TcB-TcC cocoon in TcA. Therefore, one 3D class with well-defined density of TcB-TcC was locally refined, filtered to 5 Å and used as initial template in a new 3D refinement with 1.875° initial angular sampling. The resulting map provided a clearly defined density and only one distinct orientation of TcB-TcC for both ABC(D651A) and ABC(WT).

Post-processing of the final density maps obtained by 3D refinement was performed with the PostRefiner tool of SPHIRE using a soft Gaussian mask, resulting in final average resolutions of 3.9/3.4 Å for ABC(D651A) and 3.9/3.4 Å for ABC(WT), respectively, according to FSC 0.5/0.143 (Supplementary Figures 2c, 5c). B-factors were −78.09 Å^2^ for ABC(D651A) and −61.28 Å^2^ for ABC(WT), respectively. Local FSC calculations were performed using the Local Resolution tool in SPHIRE. (Supplementary Figures 2e, 5e). The density maps were filtered according to its local resolutions using the 3D Local Filter tool in SPHIRE.

To achieve a better local resolution in the TcB-TcC part of the map of ABC(D651A), we re-boxed particles after shifting the center of the map by 70 pixels to the center of the TcB-TcC cocoon using the final projection parameters of MERIDIEN (Supplementary Figure 3a-c), and we ran MERIDIEN with the re-boxed particles. Post-processing, local FSC calculation and local filtering was performed as described previously for the full map. The overall resolution of the obtained map is 4.2/3.7 Å according to FSC 0.5/0.143 (Supplementary Figure 3d). 3D variability estimation was performed with sx3dvariability in SPHIRE. The input images were low-pass filtered to 25 Å and 20 images were used per projection.

### Atomic model building and refinement

The cryo-EM density map of ABC(D651A) was used for atomic model building and refinement. Initially, models of TcdA1 in the pore form (pdb ID 5LKH and 5LKI) and TcB-TcC (pdb ID, 6H6F) were rigid-body fitted into the density map of the holotoxin. The chains of TcA, and the TcB-TcC fusion protein were individually fitted into the density map using Imodfit (Lopéz-Blanco and Chacón, 2013) to improve the initial fit of the model. Subsequently, we modeled the atomic structure of the holotoxin using an iterative combination of manual model building in Coot (Emsley et al., 2010), Rosetta relaxation and Phenix real-space refinement (Afonine et al., 2018) without imposing symmetry. Residues 1,382-1,491 of all TcA chains were removed, as the map showed insufficient density and resolution in that regions.

After initial modeling in the full holotoxin map, the model of TcB-TcC was further improved using the shifted map (Supplementary Figure 6d,e), which allowed model building of the first residues of the ADP-ribosyltransferase of TccC3 after the mutated aspartyl protease cleavage site.

We created a fragment library of the ADP-ribosyltransferase (residues 679 – 960 of TcC) using the Robetta webserver (Kim et al., 2004) and attempted *de novo* building of the fragments into the density in the TcB channel using Rosetta. However, Rosetta *de novo* model building failed to create a reproducible model. We therefore manually built an unfolded Poly-Ala peptide in C-N translocation direction into the most well-defined density in the TcB channel. Afterwards, we added side chains at regions with sufficient density of possible peptide stretches close to the C-terminus of the ADP-ribosyltransferase and chose the best fitting model (Supplementary Figure 7b). The final models were validated by Rosetta relaxation against two independent half-maps (DiMaio et al., 2015) (Supplementary Figure 2c, 5c), EMringer (Barad et al., 2015) and MolProbity (Chen et al., 2010) (Supplementary Table 1).

### Cross-linking mass spectrometry

We incubated a 3 μM solution of TcB-TcC (WT or D651A) in 20 mM HEPES-NaOH pH 7.5, 150 mM NaCl with 3 mM disuccinimidyl dibutyric urea (DSBU, Thermo Fisher) for 30 min at 25 °C. The cross-linking reaction was stopped by adding 100 mM Tris, followed by acetone precipitation. The precipitate was dissolved in 8 M urea, 1 mM DTT and alkylated with 5.5 mM iodacetamide for 30 min at 25 °C. Afterwards, we diluted the solution 2-fold with 20 mM NH_4_HCO_3_ and digested with 1/50 molar ratio of endopeptidase LysC (Serva, MS grade) for 3 h at 25 °C. Afterwards, we diluted again 2-fold with 20 mM NH_4_HCO_3_, added 1/100 molar amount of Trypsin (Sigma, MS grade) and continued proteolysis for 12 h at 25 °C. Proteolysis was stopped by 0.1% trifluoracetic acid and the peptides were separated on a Superdex 30 increase 3.2-30 column (GE Life Science) equilibrated in 30% acetonitrile as described previously (Efremov et al., 2015). Peptides eluting between retention volumes of 1.2 and 2.0 ml were collected in 100 µl fractions, evaporated in a speedvac and subsequently analyzed via LC-MS by the mass spectrometry facility of the Max Planck Institute of Molecular Physiology. Crosslinks were evaluated using MeroX (Goetze et al., 2015) and visualized in xVis (Grimm et al., 2015).

### Bioinformatics tools

Analysis of the TcB channel diameter and constrictions was performed with ChExVis (Masood et al., 2015) and visualized in UCSF Chimera (Pettersen et al., 2004). For visualization of the Coulomb potential, TcB was protonated using H++ (Anandakrishnan et al., 2012) at pH 7.0 and an ionic strength of 150 mM. Homologs of TcdB2 were identified using Protein BLAST (Altschul et al., 1990) and sequences of TcdB2 and 6 homologs were aligned in Clustal Omega (Sievers et al., 2011). Sequence conservation of the hydrophobic and negatively charged patches in TcB and of the aspartyl protease cleavage site were analyzed with ConSurf (Ashkenazy et al., 2016) and visualized in UCSF Chimera.

## References

Afonine, P.V., Poon, B.K., Read, R.J., Sobolev, O.V., Terwilliger, T.C., Urzhumtsev, A., and Adams, P.D. (2018). Real-space refinement in Phenix for cryo-EM and crystallography.

Altschul, S.F., Gish, W., Miller, W., Myers, E.W., and Lipman, D.J. (1990). Basic local alignment search tool. J. Mol. Biol. 215, 403–410.

Anandakrishnan, R., Aguilar, B., and Onufriev, A.V. (2012). H++ 3.0: automating pK prediction and the preparation of biomolecular structures for atomistic molecular modeling and simulations. Nucleic Acids Res. 40, W537–W541.

Ashkenazy, H., Abadi, S., Martz, E., Chay, O., Mayrose, I., Pupko, T., and Ben-Tal, N. (2016). ConSurf 2016: an improved methodology to estimate and visualize evolutionary conservation in macromolecules. Nucleic Acids Res. 44, W344–W350.

Athaudaa, S.B.P., and Takahashia, K. (2002). Cleavage specificities of aspartic proteinases toward oxidized insulin B chain at different pH values. Protein Pept. Lett. 9, 289–294.

Barad, B.A., Echols, N., Wang, R.Y.-R., Cheng, Y., DiMaio, F., Adams, P.D., and Fraser, J.S. (2015). EMRinger: side chain-directed model and map validation for 3D cryo-electron microscopy. Nat. Methods 12, 943–946.

Bowen, D., Rocheleau, T.A., Blackburn, M., Andreev, O., Golubeva, E., Bhartia, R., and Ffrench-Constant, R.H. (1998). Insecticidal toxins from the bacterium photorhabdus luminescens. Science 280, 2129–2132.

Buchan, D.W.A., Minneci, F., Nugent, T.C.O., Bryson, K., and Jones, D.T. (2013). Scalable web services for the PSIPRED Protein Analysis Workbench. Nucleic Acids Res. 41, W349–W357.

Busby, J.N., Panjikar, S., Landsberg, M.J., Hurst, M.R.H., and Lott, J.S. (2013). The BC component of ABC toxins is an RHS-repeat-containing protein encapsulation device. Nature 501, 547–550.

Chen, V.B., Arendall, W.B., Headd, J.J., Keedy, D.A., Immormino, R.M., Kapral, G.J., Murray, L.W., Richardson, J.S., and Richardson, D.C. (2010). MolProbity: all-atom structure validation for macromolecular crystallography. Acta Crystallogr. D Biol. Crystallogr. 66, 12–21.

Denisov, I.G., Grinkova, Y.V., Lazarides, A.A., and Sligar, S.G. (2004). Directed self-assembly of monodisperse phospholipid bilayer Nanodiscs with controlled size. J. Am. Chem. Soc. 126, 3477–3487.

DiMaio, F., Song, Y., Li, X., Brunner, M.J., Xu, C., Conticello, V., Egelman, E., Marlovits, T., Cheng, Y., and Baker, D. (2015). Atomic-accuracy models from 4.5-Å cryo-electron microscopy data with density-guided iterative local refinement. Nat. Methods 12, 361–365.

Duchaud, E., Rusniok, C., Frangeul, L., Buchrieser, C., Givaudan, A., Taourit, S., Bocs, S., Boursaux-Eude, C., Chandler, M., Charles, J.-F., et al. (2003). The genome sequence of the entomopathogenic bacterium Photorhabdus luminescens. Nat. Biotechnol. 21, 1307–1313.

Efremov, R.G., Leitner, A., Aebersold, R., and Raunser, S. (2015). Architecture and conformational switch mechanism of the ryanodine receptor. Nature 517, 39–43.

Emsley, P., Lohkamp, B., Scott, W.G., and Cowtan, K. (2010). Features and development of Coot. Acta Crystallogr. D Biol. Crystallogr. 66, 486–501.

Ernst, K., Schmid, J., Beck, M., Hägele, M., Hohwieler, M., Hauff, P., Ückert, A.K., Anastasia, A., Fauler, M., Jank, T., et al. (2017). Hsp70 facilitates trans-membrane transport of bacterial ADP-ribosylating toxins into the cytosol of mammalian cells. Sci Rep 7, 2724.

ffrench-Constant, R.H., and Bowen, D.J. (2000). Novel insecticidal toxins from nematode-symbiotic bacteria. Cell. Mol. Life Sci. 57, 828–833.

ffrench-Constant, R.H., Waterfield, N., Burland, V., Perna, N.T., Daborn, P.J., Bowen, D., and Blattner, F.R. (2000). A genomic sample sequence of the entomopathogenic bacterium Photorhabdus luminescens W14: potential implications for virulence. Appl. Environ. Microbiol. 66, 3310–3329.

ffrench-Constant, R., and Waterfield, N. (2005). An ABC guide to the bacterial toxin complexes (Elsevier).

Gates, S.N., Yokom, A.L., Lin, J., Jackrel, M.E., Rizo, A.N., Kendsersky, N.M., Buell, C.E., Sweeny, E.A., Mack, K.L., Chuang, E., et al. (2017). Ratchet-like polypeptide translocation mechanism of the AAA+ disaggregase Hsp104. Science 357, 273–279.

Gatsogiannis, C., Lang, A.E., Meusch, D., Pfaumann, V., Hofnagel, O., Benz, R., Aktories, K., and Raunser, S. (2013). A syringe-like injection mechanism in Photorhabdus luminescens toxins. Nature 495, 520–523.

Gatsogiannis, C., Merino, F., Prumbaum, D., Roderer, D., Leidreiter, F., Meusch, D., and Raunser, S. (2016). Membrane insertion of a Tc toxin in near-atomic detail. Nat. Struct. Mol. Biol. 23, 884–890.

Gatsogiannis, C., Merino, F., Roderer, D., Balchin, D., Schubert, E., Kuhlee, A., Hayer-Hartl, M., and Raunser, S. (2018). Tc toxin activation requires unfolding and refolding of a β-propeller. Nature 563, 209–213.

Gerrard, J., Waterfield, N., Vohra, R., and ffrench-Constant, R. (2004). Human infection with Photorhabdus asymbiotica: An emerging bacterial pathogen. Microbes and Infection 6, 229–237.

Goetze, M., Pettelkau, J., Fritzsche, R., Ihling, C.H., Schaefer, M., and Sinz, A. (2015). Automated Assignment of MS/MS Cleavable Cross-Links in Protein 3D-Structure Analysis. J. Am. Soc. Mass Spectrom. 26, 83–97.

Grimm, M., Zimniak, T., Kahraman, A., and Herzog, F. (2015). xVis: a web server for the schematic visualization and interpretation of crosslink-derived spatial restraints. Nucleic Acids Res. 43, W362–W369.

Hessa, T., Kim, H., Bihlmaier, K., Lundin, C., Boekel, J., Andersson, H., Nilsson, I., White, S.H., and Heijne von, G. (2005). Recognition of transmembrane helices by the endoplasmic reticulum translocon. Nature 433, 377–381.

Jackson, V.A., Meijer, D.H., Carrasquero, M., van Bezouwen, L.S., Lowe, E.D., Kleanthous, C., Janssen, B.J.C., and Seiradake, E. (2018). Structures of Teneurin adhesion receptors reveal an ancient fold for cell-cell interaction. Nat Commun 9, 1079.

Jiang, J., Pentelute, B.L., Collier, R.J., and Hong Zhou, Z. (2015). Atomic structure of anthrax protective antigen pore elucidates toxin translocation. Nature 521, 545–549.

Kim, D.E., Chivian, D., and Baker, D. (2004). Protein structure prediction and analysis using the Robetta server. Nucleic Acids Res. 32, W526–W531.

Krantz, B.A., Melnyk, R.A., Zhang, S., Juris, S.J., Lacy, D.B., Wu, Z., Finkelstein, A., and Collier, R.J. (2005). A phenylalanine clamp catalyzes protein translocation through the anthrax toxin pore. Science 309, 777–781.

Lang, A.E., Ernst, K., Lee, H., Papatheodorou, P., Schwan, C., Barth, H., and Aktories, K. (2014). The chaperone Hsp90 and PPIases of the cyclophilin and FKBP families facilitate membrane translocation of Photorhabdus luminescensADP-ribosyltransferases. Cell. Microbiol. 16, 490–503.

Lang, A.E., Schmidt, G., Schlosser, A., Hey, T.D., Larrinua, I.M., Sheets, J.J., Mannherz, H.G., and Aktories, K. (2010). Photorhabdus luminescens toxins ADP-ribosylate actin and RhoA to force actin clustering. Science 327, 1139–1142.

Li, J., Shalev-Benami, M., Sando, R., Jiang, X., Kibrom, A., Wang, J., Leon, K., Katanski, C., Nazarko, O., Lu, Y.C., et al. (2018). Structural Basis for Teneurin Function in Circuit-Wiring: A Toxin Motif at the Synapse. Cell 173, 735–748.e15.

Lopéz-Blanco, J.R., and Chacón, P. (2013). iMODFIT: efficient and robust flexible fitting based on vibrational analysis in internal coordinates. J. Struct. Biol. 184, 261–270.

Masood, T.B., Sandhya, S., Chandra, N., and Natarajan, V. (2015). CHEXVIS: a tool for molecular channel extraction and visualization. BMC Bioinformatics 16, 119.

Meusch, D., Gatsogiannis, C., Efremov, R.G., Lang, A.E., Hofnagel, O., Vetter, I.R., Aktories, K., and Raunser, S. (2014). Mechanism of Tc toxin action revealed in molecular detail. Nature 508, 61–65.

Morán Luengo, T., Kityk, R., Mayer, M.P., and Rüdiger, S.G.D. (2018). Hsp90 Breaks the Deadlock of the Hsp70 Chaperone System. Mol. Cell 70, 545–552.e549.

Moriya, T., Saur, M., Stabrin, M., Merino, F., Voicu, H., Huang, Z., Penczek, P.A., Raunser, S., and Gatsogiannis, C. (2017). High-resolution Single Particle Analysis from Electron Cryo-microscopy Images Using SPHIRE. J Vis Exp 2017, e55448–e55448.

Penczek, P.A., Fang, J., Li, X., Cheng, Y., Loerke, J., and Spahn, C.M.T. (2014). CTER-rapid estimation of CTF parameters with error assessment. Ultramicroscopy 140, 9–19.

Pettersen, E.F., Goddard, T.D., Huang, C.C., Couch, G.S., Greenblatt, D.M., Meng, E.C., and Ferrin, T.E. (2004). UCSF Chimera--a visualization system for exploratory research and analysis. J Comput Chem 25, 1605–1612.

Puchades, C., Rampello, A.J., Shin, M., Giuliano, C.J., Wiseman, R.L., Glynn, S.E., and Lander, G.C. (2017). Structure of the mitochondrial inner membrane AAA+ protease YME1 gives insight into substrate processing. Science 358, eaao0464.

Sergeant, M., Jarrett, P., Ousley, M., and Morgan, J.A.W. (2003). Interactions of insecticidal toxin gene products from Xenorhabdus nematophilus PMFI296. Appl. Environ. Microbiol. 69, 3344–3349.

Sievers, F., Wilm, A., Dineen, D., Gibson, T.J., Karplus, K., Li, W., Lopez, R., McWilliam, H., Remmert, M., Söding, J., et al. (2011). Fast, scalable generation of high-quality protein multiple sequence alignments using Clustal Omega. Mol. Syst. Biol. 7, 539–539.

Tabashnik, B.E., Brévault, T., and Carrière, Y. (2013). Insect resistance to Bt crops: lessons from the first billion acres. Nat. Biotechnol. 31, 510–521.

Tennant, S.M., Skinner, N.A., Joe, A., and Robins-Browne, R.M. (2005). Homologues of insecticidal toxin complex genes in Yersinia enterocolitica biotype 1A and their contribution to virulence. Infect. Immun. 73, 6860–6867.

Wagner, T., Merino, F., Stabrin, M., Moriya, T., Gatsogiannis, C., and Raunser, S. (2018). SPHIRE-crYOLO: A fast and well-centering automated particle picker for cryo-EM. bioRxiv 356584.

Waterfield, N.R., Bowen, D.J., Fetherston, J.D., Perry, R.D., and ffrench-Constant, R.H. (2001). The tc genes of Photorhabdus: a growing family. Trends Microbiol. 9, 185–191.

Waterfield, N., Hares, M., Hinchliffe, S., Wren, B., and ffrench-Constant, R. (2007). The insect toxin complex of Yersinia. Adv. Exp. Med. Biol. 603, 247–257.

Yang, G., and Waterfield, N.R. (2013). The role of TcdB and TccC subunits in secretion of the Photorhabdus Tcd toxin complex. PLoS Pathog. 9, e1003644.

Yang, Z., Fang, J., Chittuluru, J., Asturias, F.J., and Penczek, P.A. (2012). Iterative stable alignment and clustering of 2D transmission electron microscope images. Structure 20, 237–247.

Zhang, K., Li, M., and Sun, F. (2011). Gautomatch: an efficient and convenient gpu-based automatic particle selection program.

Zheng, S.Q., Palovcak, E., Armache, J.-P., Verba, K.A., Cheng, Y., and Agard, D.A. (2017). MotionCor2: anisotropic correction of beam-induced motion for improved cryo-electron microscopy. Nat. Methods 14, 331–332.

